# Transcriptional Regulation of Synthetic Polymer Networks

**DOI:** 10.1101/2021.10.17.464678

**Authors:** Austin J. Graham, Christopher M. Dundas, Gina Partipilo, Ismar E. Miniel Mahfoud, Thomas FitzSimons, Rebecca Rinehart, Darian Chiu, Avery E. Tyndall, Adrianne M. Rosales, Benjamin K. Keitz

## Abstract

Individual cells direct non-equilibrium processes through coordinated signal transduction and gene expression, allowing for dynamic control over multicellular, system-wide behavior. This behavior extends to remodeling the extracellular polymer matrix that encases biofilms and tissues, where constituent cells dictate spatiotemporal network properties including stiffness, pattern formation, and transport properties. The majority of synthetic polymer networks cannot recreate these phenomena due to their lack of autonomous centralized actuators (*i*.*e*., cells). In addition, non-living polymer networks that perform computation are generally restricted to a few inputs (*e*.*g*., light, pH, enzymes), limiting the logical complexity available to a single network chemistry. Toward synergizing the advantages of living and synthetic systems, engineered living materials leverage genetic and metabolic programming to establish control over material-wide properties. Here we demonstrate that a bacterial metal respiration mechanism, extracellular electron transfer (EET), can control metal-catalyzed radical cross-linking of polymer networks. Linking metabolic electron flux to a synthetic redox catalyst allows dynamic, tunable, and predictable control over material formation and bulk polymer network mechanics using genetic circuits. By programming key EET genes with transcriptional Boolean logic, we rationally design computational networks that sense-and-respond to multiple inputs in biological contexts. Finally, we capitalize on the wide reactivity of EET and redox catalyses to predictably control another class of living synthetic materials using copper(I) alkyne-azide cycloaddition click chemistry. Our results demonstrate the utility of EET as a bridge for controlling abiotic materials and how the design rules of synthetic biology can be applied to emulate physiological behavior in polymer networks.

## Main

Living polymer networks, including tissues and biofilms, actively respond to complex combinations of environmental inputs, leading to phenomena such as differentiation, regeneration, and development^1–3^. In these systems, the bidirectional flow of chemical^4^, mechanical^5–7^, and electrical^8,9^ information between constituent cells and their extracellular matrix informs micro- and macroscopic morphology and function. These dynamic capabilities are made possible through distributed sensing, amplification, and actuation machinery controlled by genetic regulatory networks^10,11^. In contrast, synthetic polymer networks generally lack similar mechanisms for active computation. Efforts to program non-living polymer networks with biomimetic behavior have received significant attention^12–18^, but network transformations using purely chemical approaches typically require large input signals and multiple orthogonal chemistries, limiting their ability to evolve beyond relatively simple operations^19–23^. Addressing the limitations of abiotic materials, some implementations of engineered living materials capitalize on the programmability of resident cells to control material synthesis and system-wide properties^24–27^. However, these strategies generally rely on materials natively produced by the host organism, potentially narrowing their application. Augmenting the functional advantages of synthetic materials with the sensing and computing capabilities of biological systems could advance several fields, including tissue engineering^28,29^, additive manufacturing^30^, and artificial signaling^31,32^.

Here we describe synthetic polymer networks (*i*.*e*., hydrogels) that dynamically cross-link in response to multi-input computations performed by resident bacteria. Our approach coopts extracellular electron transfer (EET)^33^, a respiratory mechanism by which microbes transport electrons across their membrane to reduce a variety of soluble metal species, to connect cellular biochemical reaction networks to the activity of exogenous metal catalysts^34,35^. Leveraging the transcriptional programmability of EET genes in the model electroactive bacterium *Shewanella oneidensis*, we deploy engineered bacteria as distributed computing and actuating elements within a solution-phase network precursor. Upon sensing dilute chemical stimuli, *S. oneidensis* strains transcriptionally regulate expression of EET pathways and subsequently activate a metal radical polymerization catalyst, which cross-links the network. Using this platform, we first show that cross-linking can be dynamically coupled to gene expression to predictably control polymer network mechanics through transcriptional regulation. We then expand the logical complexity of these networks by engineering two-input genetic Boolean circuits (OR, NOR, AND, and NAND) for controlling EET and resultant gel mechanics. Finally, capitalizing on the modularity of EET, we demonstrate that genetic logic can control another well-known cross-linking chemistry, copper(I)-catalyzed alkyne-azide cycloaddition (CuAAC). By coupling biological computation to the activity of metal cross-linking catalysts, we showcase EET’s potential as a universal interface for programming polymer network properties and forward-engineering living materials using genetic circuits^36,37^.

### Relative Gene Expression Predicts Network Mechanics

We initially hypothesized that polymer network synthesis could be connected to stimuli-responsive genetic circuity designed to control EET gene expression. For example, we previously demonstrated that a transcriptional circuit regulating a single EET gene, *mtrC*, in *S. oneidensis* could predictably control the mechanical properties of cross-linked hydrogels formed via radical polymerization^35^. In this system, EET to a copper redox catalyst controlled the cross-linking rate and storage modulus of methacrylated macromers through atom-transfer radical polymerization^38^. To simplify *mtrC* circuit assessment, we previously separated the dynamics of EET gene expression and polymerization by growing bacteria in inducer before applying them to cross-linking reactions at stationary phase. However, biomimetic materials should ideally sense environmental signals in real-time and actuate physical changes accordingly. We therefore coupled bacterial sensing, gene expression, and growth to cross-linking, such that hydrogels formed dynamically upon inoculation with naive *S. oneidensis* cells at low initial density with appropriate inducers (Figure 1a, Extended Data Fig. 1, “dynamic” cross-linking). Methacrylate-functionalized polyethylene glycol networks formed using previously induced or naive cells exhibited similar regulation of storage modulus when using a common regulatory motif, the LacI-*P*_*tacsymO*_ regulator-promoter pair, to control *mtrC* expression in a Δ*mtrC*Δ*omcA*Δ*mtrF* knockout (JG596)^39^. This manifested as characteristic response functions and similar fold-change between ON/OFF states in response to inducer (isopropyl-β-D-thiogalactopyranoside, IPTG), despite the additional requirement for coordinated growth, protein expression, and cross-linking under dynamic conditions. Cells remained viable and metabolically active for up to a week in the gels (Extended Data Fig. 2), and strains harboring empty vector controls formed weaker gels under all conditions. Alternative EET machinery (*mtrA, cymA*) could also be transcriptionally regulated toward dynamic network cross-linking, highlighting the programmability of EET pathways in *S. oneidensis* for material synthesis. Because the *mtrC* circuits exhibited high dynamic range and the resultant protein is the direct actuator of catalyst reduction, we opted to control *mtrC* expression in subsequent genetic circuits.

**Figure 1.**
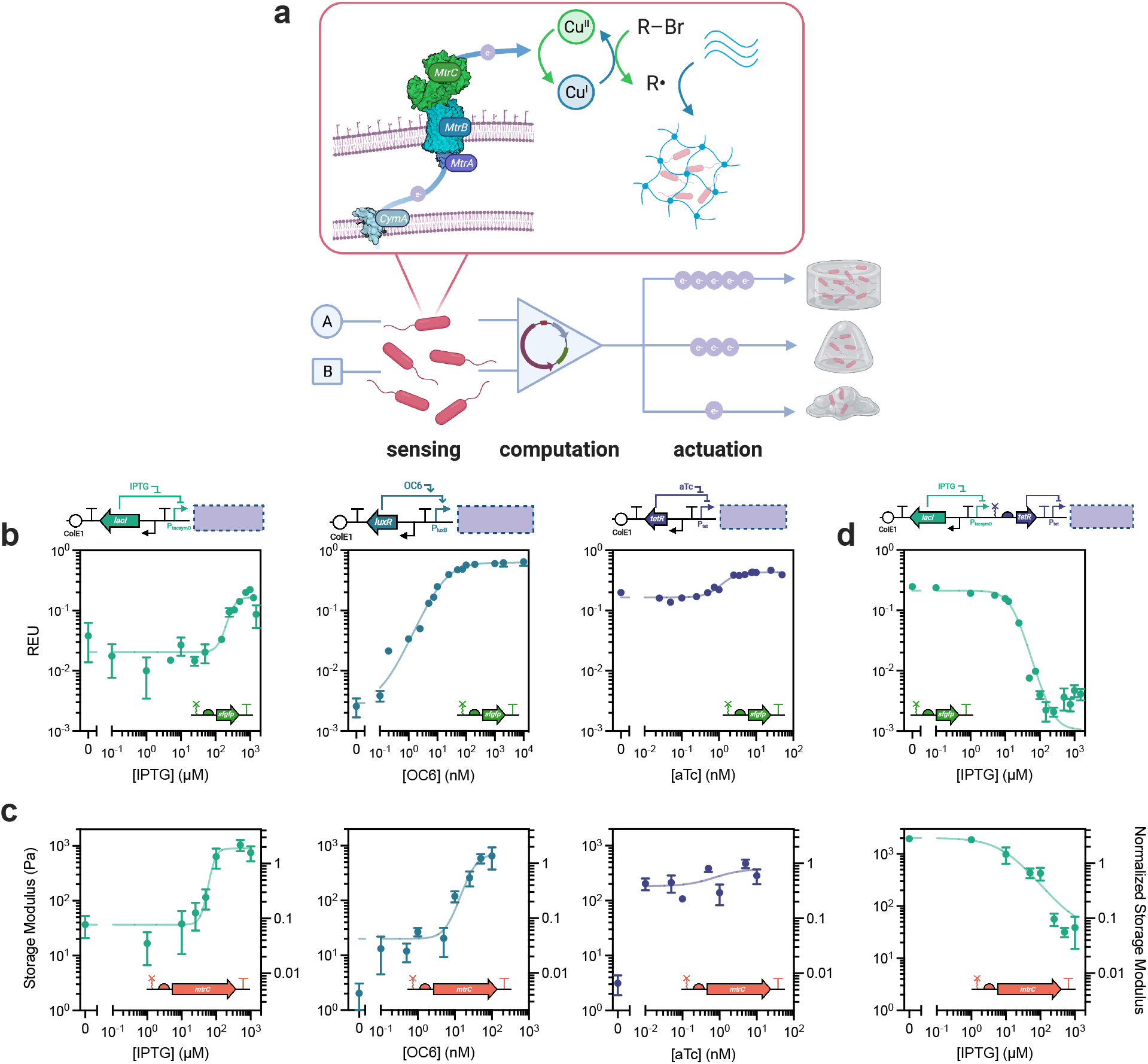
Various transcriptional architectures enable quantitative control over gene expression and resultant polymer network mechanics. **a**, Schematic illustrating computational polymer networks actuated by transcriptional logic. *S. oneidensis* acts as a distributed computing element within a network precursor solution, and input signals activate or deactivate an extracellular electron transfer pathway. This pathway is comprised of various cytochrome proteins, which control the redox cycle of a catalytic metal. Upon reduction, the metal catalyst powers an orthogonal chemical reaction such as atom-transfer radical polymerization. The result is a synthetic polymer network whose properties are coupled to biological computation. **b**, Relative Expression Units (REU) measured as a function of inducer concentration show characteristic transcriptional regulation over a variety of dynamic ranges for multiple traditional Buffer gate architectures expressing *sfgfp*. The corresponding circuit architecture for each response function is shown above as a cartoon, with representations for origins of replication, terminators, promoters, ribozymes, RBS, and genes; the cognate inducing molecule is shown as well. **c**, Storage moduli of methacrylated PEG polymer networks dynamically cross-linked using the same architectures as in **b**, but controlling *mtrC* expression, measured 18 h after inoculation. The response functions for a physical material property mirror those predicted by relative gene expression. **d**, REU and storage modulus response functions as in **b** and **c**, but for a NOT gate architecture demonstrating transcriptional turn-off, measured 24 h after inoculation. Data shown are mean ± SEM of *n* = 3 biological replicates.

Since these circuits were optimized for responsiveness to IPTG^40^, we anticipated that changes to the inducer-sensing regulator or overall circuit topology could enable new computation, but potentially cause variable or unintended circuit functioning (*e*.*g*., leaky *mtrC* expression resulting in high background EET). In similar transcriptional circuits that regulate RNA polymerase flux, parameterization of circuit outputs using fluorescent reporter Relative Expression Units (REUs) facilitated circuit debugging and optimization^36,37^. As REU measurements can enable forward-engineering of gene expression and design of complex cellular logic, we leveraged this metrology to probe how new *mtrC* circuits affected hydrogel formation and developed design rules for programming EET-driven materials synthesis. Specifically, we parameterized various single-regulator transcriptional circuits to activate gene expression and EET activity (*i*.*e*., Buffer gates). We constructed plasmids containing the regulator-promoter pairs LacI-*P*_*tacsymO*_, LuxR-*P*_*Lux*_, and TetR-*P*_*Tet*_ to drive transcription of either *sfgfp* or *mtrC* in response to their cognate inducers (IPTG; 3-oxohexanoyl-homoserine lactone, OC6; and anhydrotetracycline, aTc, respectively). Using *S. oneidensis* strains transformed with *sfgfp* circuits, we initially assessed the transcriptional control of each Buffer gate by measuring OD_600_-normalized fluorescence after overnight growth in inducer-containing media. REU values were determined by normalizing measurements to fluorescence from an *S. oneidensis* strain carrying a constitutive *sfgfp* plasmid. All three Buffer gates exhibited characteristic ‘turn-on’ response functions, where REU values increased in the presence of increasing inducer concentration (Figure 1b). Overall, each Buffer gate exhibited varying REU ON/OFF states and dynamic ranges, and these trends were validated by fitting to a sigmoidal gene expression model.

Next, we assessed these Buffer gate architectures for tailored *mtrC* expression and subsequent gel formation using engineered *S. oneidensis* strains. In contrast to *sfgfp* fluorescence measurements, which are static, *mtrC* expression and cross-linking are both dynamic processes. Thus, polymer network mechanics will change with time, even under conditions approximating steady-state *mtrC* expression. We therefore initially selected cross-linking reaction times that optimized ON/OFF mechanical differences between gels formed using induced and uninduced strains. Each *mtrC* harboring strain was simultaneously inoculated and induced into gel precursor solutions and storage modulus was measured after 18 h, which was found to emphasize transcriptional differences (Figure 1c). Response functions for dynamically cross-linked networks closely mirrored the response for each REU-parameterized gate, indicating similar transcriptional regulation and validating forward-engineering of material properties from established gene expression models. In all cases, gels did not form in the presence of induced strains harboring an empty vector control. Notably, we found that gel stiffness increased concomitantly with inducer concentration across all Buffer gate circuits, except when the inducer/circuit combination yielded REU values above *ca*. 0.2–0.3. This was especially noticeable with the TetR-regulated Buffer gate where, across all aTc concentrations, the transcriptional output was above this range. For the LacI and LuxR Buffer gates that operated below this REU bound, gel mechanics fit well to gene expression models and exhibited higher dynamic ranges, spanning a roughly 1000-fold range in mechanical properties. Together, these results demonstrate that well-characterized inducers and regulators can activate *mtrC*-driven gel formation and highlight useful design rules within our platform for establishing transcriptional control over network formation using MtrC.

To enable more complex logical computation, EET flux and associated polymerization activity must also turn off in response to transcriptional regulation. However, when deactivating gene expression, growth and cell division (minutes–hours) are required to dilute proteins whose turnover rates exceed the timescale of cell division, as is the case with MtrC (half-life *ca*. 16 h)^41,42^. For dynamic cross-linking, this dilution must also out-compete polymerization kinetics (*ca*. minutes after catalyst activation). To address these challenges, we examined a previously designed two-regulator NOT gate, which represses *mtrC* transcription and EET activity in response to IPTG, for hydrogel formation^40^. After optimizing reaction time, we found that networks cross-linked by *S. oneidensis* carrying this NOT gate formed weaker networks in response to increasing IPTG concentration, confirming dynamic repression of EET and resultant material properties (Figure 1d). The response function of this NOT gate also reflected that EET could be turned off only if REU could be tuned below 0.2–0.3. Overall, longer reaction times (24 h) enabled coordinated transcriptional repression of polymerization activity, confirming that dynamic *mtrC* regulation could also attenuate cross-linking activity using genetic circuits.

### Boolean Logic Programs Polymer Network Dynamics

Successful hydrogel formation using different Buffer and NOT gate architectures regulating EET suggested that *S. oneidensis* could be engineered to perform more complex logical operations during polymer network synthesis. To demonstrate this, we engineered genetic two-input Boolean logic based on nested repressor architectures to control *mtrC* expression and associated cross-linking (Figure 2)^36,37,43,44^. Each gate was designed with a sensing block using combinations of common small molecule inducers: OC6 and aTc sensors for OR and NOR gates; IPTG and aTc sensors for AND gates; and IPTG and OC6 sensors for NAND gates. These sensor blocks control activation and deactivation of various repressor proteins, which coordinate to yield the desired gate-dependent transcriptional output. We first functionally validated each circuit in *S. oneidensis* using fluorescent gene expression (Figure 2). Logical output was a function of repressor architecture and inducer concentration, and 2D heat maps reflected expected gradients in REU (Extended Data Fig. 3). Thus, we confirmed the expected truth tables for each genetic Boolean architecture and functionally validated many repressor proteins not previously used in *S. oneidensis*.

**Figure 2.**
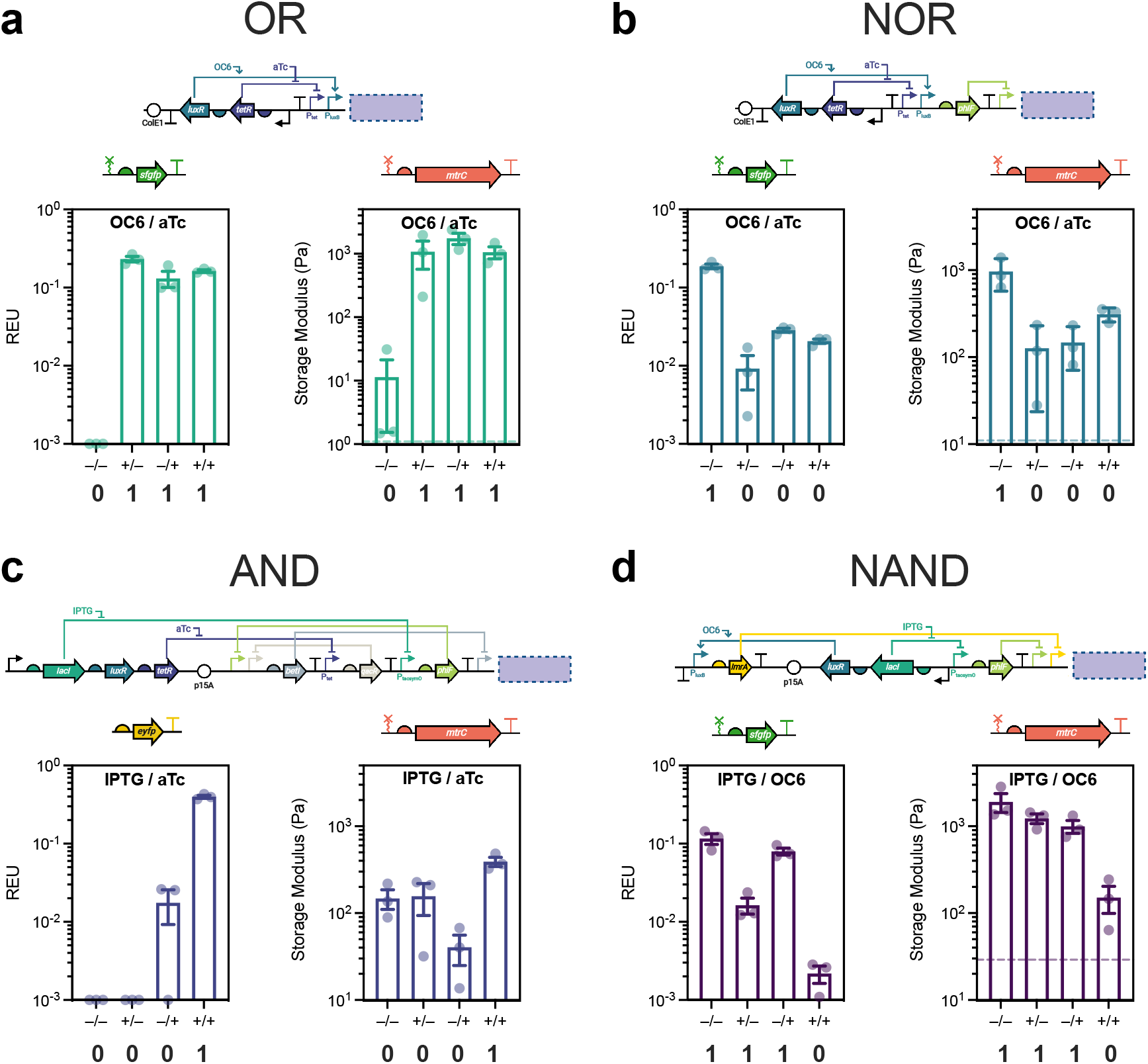
Genetic Boolean logic enables polymer network computation via living cellular actuators. Relative gene expression (REU) and polymer network mechanics (storage modulus) using *S. oneidensis* strains harboring transcriptional Boolean logic circuits controlling fluorescent gene (left) or *mtrC* (right) expression. The logical architectures span **a**, OR, **b**, NOR, **c**, AND, and **d**, NAND using nested repressors that coordinate in response to sensor blocks. The repressors are activated or deactivated in response to their cognate inducing molecules (100 μM IPTG, 100 nM OC6, 10 nM aTc, and/or their corresponding solvents). In all cases, storage modulus was measured 24 h after inoculation. The expected truth tables are represented below each circuit (“0”, OFF; “1”, ON). Each plasmid architecture is shown as a cartoon above the corresponding response function, with representations as in Figure 1. In general, each strain appropriately responds to combinatorial inputs by increasing/decreasing fluorescence or storage modulus in response to fluorescent gene or *mtrC* activation/deactivation. Where applicable, dashed lines represent average storage modulus of gels formed using a fluorescent vector control. Data shown are mean ± SEM of *n* = 3 biological replicates.

We then placed *mtrC* under genetic Boolean regulation and applied these engineered strains toward polymer network cross-linking. Gates were tested for both stationary phase and dynamic cross-linking. In all cases, resultant storage modulus followed the expected truth table for each logic operation (Figure 2, Extended Data Fig. 4). The OR gate, which was the simplest two-input architecture, exhibited the greatest dynamic range in storage modulus (∼100-fold). NOR, NAND, and AND, which are increasingly complex architectures, exhibited dynamic ranges of ∼3 to 10-fold. The inducer-dependent outputs from the AND and NAND gates matched well with previous reports that used these specific architectures^44^, including greater overall transcription in the NAND gate and decreased dynamic range in the AND gate. Strains harboring each circuit showed no measurable growth defect (Extended Data Fig. 5), supporting the role of EET-driven gel formation over unintended circuit effects. Gate-based regulation of EET was also validated in separate Fe(III) reduction assays (Extended Data Fig. 6). Finally, gels formed in the presence of control strains harboring induced fluorescent vectors were significantly weaker than *mtrC* expressing strains or did not form at all. Overall, logical computation in *S. oneidensis* using these 2-input gates rationally tuned polymer network mechanics within biologically relevant windows, highlighting the utility of transcriptional regulation for programmable material computation through EET.

### Extracellular Electron Transfer Actuates an Alternative Cross-linking Chemistry

A significant advantage of EET is its potential to influence other metal-catalyzed reactions. As a specific example, we predicted that *S. oneidensis* could transcriptionally regulate copper(I)-catalyzed alkyne-azide cycloaddition (CuAAC) click chemistry using the Mtr pathway^45^. CuAAC is a ubiquitous bioorthogonal cross-linking reaction that accesses unique polymer network structures compared to radical polymerizations by virtue of a step-growth mechanism^38,46^. Using 4-arm alkyne- and azide-functionalized PEG macromers, we first demonstrated that CuAAC cross-linking was unique to EET-capable *S. oneidensis* strains and resultant hydrogel mechanics were genetically encoded (Figure 3a). After establishing this link, we investigated transcriptional control using Buffer and NOT gate architectures controlling *mtrC* expression. In each case, regulation of EET activity in response to an inducer enabled dynamic control over CuAAC hydrogel mechanics after 12 h of cross-linking (Figure 3b, c). The response function for each gate exhibited expected behavior within a biologically relevant window, with increased dynamic range relative to EET-controlled radical polymerization. In addition, the ON/OFF behavior of each gate could be modulated by changing the Cu ligand (Extended Data Fig. 7), which influences EET-driven CuAAC activity^45^.

**Figure 3.**
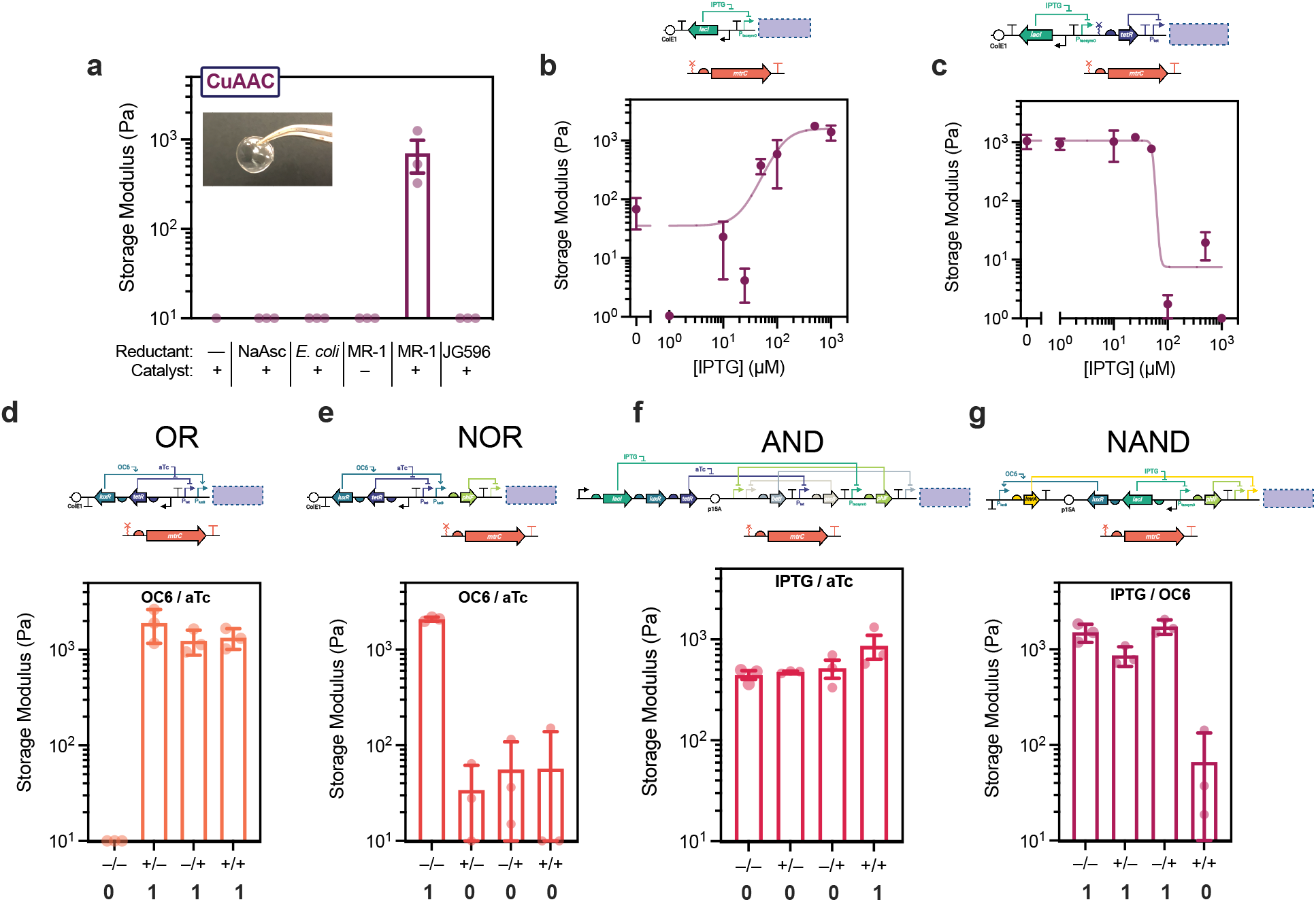
New living material chemistries enabled by EET-driven CuAAC cross-linking. **a**, Storage moduli of CuAAC cross-linked polymer networks using a variety of chemical or biological reductants and a Cu catalyst. Gels do not form without EET activity via extracellular cytochromes as evidenced by lack of gelation using *E. coli* or *S. oneidensis* JG596, demonstrating genetic control over the reaction. Inset is a representative picture of a CuAAC cross-linked hydrogel. **b**–**c**, CuAAC cross-linked polymer networks can be transcriptionally regulated via EET using **b**, Buffer and **c**, NOT gate architectures. **d**–**g** Genetic Boolean logic enables new network chemistries, using EET-driven CuAAC cross-linking to control *mtrC* expression for **d**, OR, **e**, NOR, **f**, AND, and **g**, NAND architectures. In all cases, storage modulus was measured 12 h after inoculation. The expected truth tables are represented below each circuit (“0”, OFF; “1”, ON). Each plasmid architecture is shown as a cartoon above the corresponding response function, with representations as in Figure 1. Data shown are mean ± SEM of *n* = 3 biological replicates.

Given that *mtrC* expression could predictably control CuAAC activity, we expected that this cross-linking mechanism could also mechanically regulate hydrogels using our genetic Boolean logic constructs. Indeed, gels formed dynamically in response to appropriate environmental inputs and followed the expected logic (Figure 3d–g). Network transformations generally exhibited improved dynamic range compared to radical polymerization, with accelerated cross-linking kinetics (12 h). The only exception was the AND gate, being the most complex regulatory architecture, where transcriptional differences were possibly muted by rapid CuAAC kinetics^45,47^. In all cases, gels did not form in the presence of induced vector controls. Ultimately, successful functioning of these circuits further demonstrates EET’s programmability and highlights its potential to serve as a modular genetic actuator for catalyzing diverse synthetic chemistries. The generality of redox-powered cross-linking could be applied to create many new living material platforms that respond to transcriptional regulation.

## Discussion

By linking metabolic function to a synthetic catalyst, we created a generalizable platform for living material synthesis that capitalizes on the diverse computational power of transcriptional logic. Specifically, we developed dynamic cross-linking reactions where cell growth and sensing were coupled to polymer network formation, emulating natural processes such as biofilm and tissue formation. This led to measurable gradients in gene expression (REU) and living material properties (storage modulus) based on real-time molecular sensing. Despite the variety of timescales associated with transcription, protein production/transport, EET to the metal catalyst, and polymerization, we observed significant control over hydrogel storage modulus, even for complex transcriptional architectures. Our observation that storage moduli fit well to characteristic gene expression models is supported by theory in that during exponential growth, protein concentrations approach steady-state^48,49^. Thus, actively growing cells should reflect predictable changes in induction and EET activity. We also note that our parametrization of EET gene expression used REU measurements taken at stationary phase; these values likely differ during early exponential growth in gels. Nonetheless, our REU metrology still enabled predictable design rules for hydrogel formation, with accuracy that may be improved by kinetic fluorescence measurements^49^. While synthetic biologists have established numerous paradigms for programming living systems, including computation using genetic logic, the majority have focused on fluorescence as an output^37,49–54^. By applying these concepts toward adapting circuits for controlling EET gene expression and subsequent catalyst activation, we initiated development of parallel design rules for living materials, where mechanical properties of polymer networks depend on transcriptional output.

In addition to cell growth and protein expression, developing living materials requires consideration of enzymatic and chemical kinetics. Our platform provides a foundation for merging and probing fundamentals from these fields toward systems biology-inspired material design. However, this complexity leads to non-idealities that do not exist in purely chemical systems, including leaky gene expression, metabolic burden, and non-specific catalyst activation. For example, cross-linking in our system can occur independently of transcriptional regulation via background radical production or adventitious catalyst reduction. Fortunately, we found that even relatively complex transcriptional initiation could still out-compete these processes to yield the predicted output. We did observe differences in gate performance and dynamic range depending on circuit identity. Specifically, the AND gate, which is the most convoluted architecture, exhibited diminished dynamic range compared to the other gates, while still generally performing as predicted. Our results are consistent with the observation that nested transcriptional architectures are generally not favored by natural systems to control time-dependent processes requiring fast dynamics^55^. However, a significant advantage of our system is the direct genetic link between EET and catalyst activity, implying that cross-linkable materials are theoretically controllable using any regulation of gene expression, including CRISPR^50^, riboswitches^51,52^, or integrases^53,54^.

Employing live cellular actuators in synthetic polymer networks allows the cross-linking chemistry to remain independent of the input signal and logical architecture. This orthogonality vastly broadens the scope of inputs and computations that can regulate a cross-linking event, including signals such as specific DNA/RNA sequences^50,51^, clinical biomarkers^56,57^, or environmental contaminants^58^. Applying recent advancements in genetic circuit standardization, such as gate matching^37^, could further tune network performance and sensitivity to tailor the cross-linking response from signal concentration-dependent (analog) to binary (digital). Relative to chemical systems, our platform responds to very low input magnitudes (pM–nM) due to the sensitivity of biological machinery, but actuates macroscopic material transformations that span a ∼1000-fold physiologically relevant mechanical range. Additionally, EET-controlled radical polymerization and CuAAC demonstrate significant expansions of the chemical reaction space available to microbes, but remain compatible with other biomaterials, such as tissue engineering scaffolds^19^, cellulose^26^, and curli fibers^27^. The two cross-linking chemistries presented here greatly vary in mechanism, yet both could be genetically and transcriptionally regulated. Biological control over these orthogonal chemistries using identical genetic machinery also suggests that EET can serve as a modular actuator for other synthetic redox reactions, providing a platform for harnessing advances in biomaterial engineering, organometallic chemistry, and synthetic biology^24,28,59^.

Researchers from numerous fields increasingly seek to capitalize on biology’s computational diversity to enhance traditionally non-living systems. Synthetic biology has evolved into a standardized discipline of user-defined sensors, regulators, and outputs. Living materials can leverage these tools, but have predominately relied on genetic control of host-synthesized materials or chemistries with limited substrate scope. Here we applied this foundation toward the development of living synthetic materials that intelligently respond to molecular inputs through the biological interface of EET. This direct genetic link between microbial physiology and redox chemistry enabled predictable transcriptional regulation over synthetic polymer networks. Our design’s programmability holds promise for intervention-less and dynamic applications, where engineered microbes are incorporated into tissue architectures, biosensors, soft actuators, or additive manufacturing platforms. Even relatively small changes in local micromechanics are critical in living systems, and the ability to transcriptionally tune within this window may open new biomimetic opportunities. Overall, our work provides the foundation for applying the transcriptional regulatory motifs found in patterning, embryogenesis, and tissue formation toward the control of synthetic polymer networks.

## Acknowledgments

Base plasmids for the AND and NAND circuits were generously provided by the Voigt Lab via Addgene (#49375, #49376, #49377). This research was financially supported by the Welch Foundation (Grant F-1929, B.K.K.), the National Institutes of Health under award number R35GM133640 (B.K.K.), an NSF CAREER award (1944334, B.K.K.), and the Air Force Office of Scientific Research under award number FA9550-20-1-0088 (B.K.K.). A.J.G. and G.P. were supported through National Science Foundation Graduate Research Fellowships (Program Award No. DGE-1610403). The authors acknowledge use of shared research facilities supported in part by the Texas Materials Institute, the Center for Dynamics and Control of Materials: an NSF MRSEC (DMR-1720595), and the NSF National Nanotechnology Coordinated Infrastructure (ECCS-1542159). A.M.R. gratefully acknowledges a Career Award at the Scientific Interface (#1015895) from the Burroughs Wellcome Fund. We gratefully acknowledge the use of facilities within the core microscopy lab of the Institute for Cellular and Molecular Biology, University of Texas at Austin. NMR spectra were collected on a Bruker Avance III HD 400 funded by the NSF (Award CHE 1626211). Schematics were created using BioRender.com.

## Author Contributions

A.J.G., C.M.D., G.P., and B.K.K. conceived the project and designed research. A.J.G., G.P., and D.C. performed cross-linking experiments and rheological analysis. A.J.G., C.M.D., I.E.M., R.R., and A.E.T. performed cloning and circuit characterization by growth, fluorescence, and iron reduction assays. T.F. synthesized alkyne-functionalized PEG and provided reagents. A.M.R. and B.K.K. supervised research. A.J.G., C.M.D., and B.K.K. wrote the paper with input from all authors.

## Materials & Correspondence

Request for materials and correspondence should be addressed to B.K.K.

## Data Availability

Experimental data supporting the findings of this study will be available through the Texas Data Repository.

---

**Extended Data Fig. 1.**
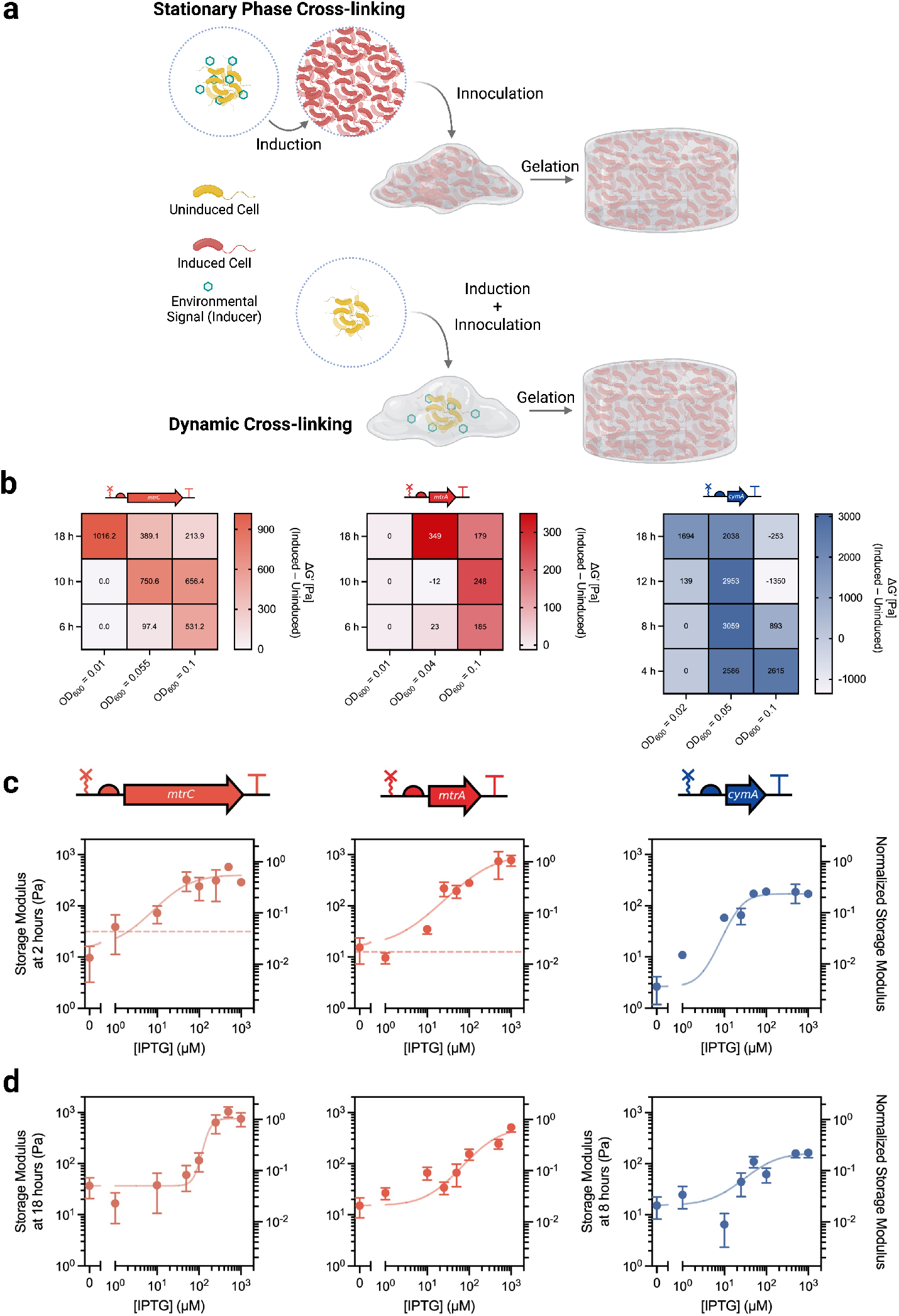
Dynamic cross-linking couples sensing, computation, and actuation in a synthetic material. **a**, Schematic illustrating stationary phase vs. dynamic cross-linking. **b**, Representative results of cross-linking condition scan to identify optimal transcriptional control over dynamic networks using *mtrC, mtrA*, or *cymA*. A value of 0 represents no measurable gel formation. Parameters such as Cu-ligand and radical initiator concentration were also explored. **c**–**d**, Cross-linking can be transcriptionally regulated using the LacI-*P*_*tacsymO*_ regulator-promoter pair controlling *mtrC, mtrA*, or *cymA* expression under **c**, stationary phase or **d**, dynamic conditions. Data are fit to an activating gene expression model and right axis is normalized to storage modulus of gels formed using wild-type *S. oneidensis* harboring an empty vector. Dashed lines represent gel mechanics using corresponding knockout strains harboring an empty vector; if no line is shown, gels did not form. Data shown are mean ± SEM of *n* = 3 biological replicates.

**Extended Data Fig. 2.**
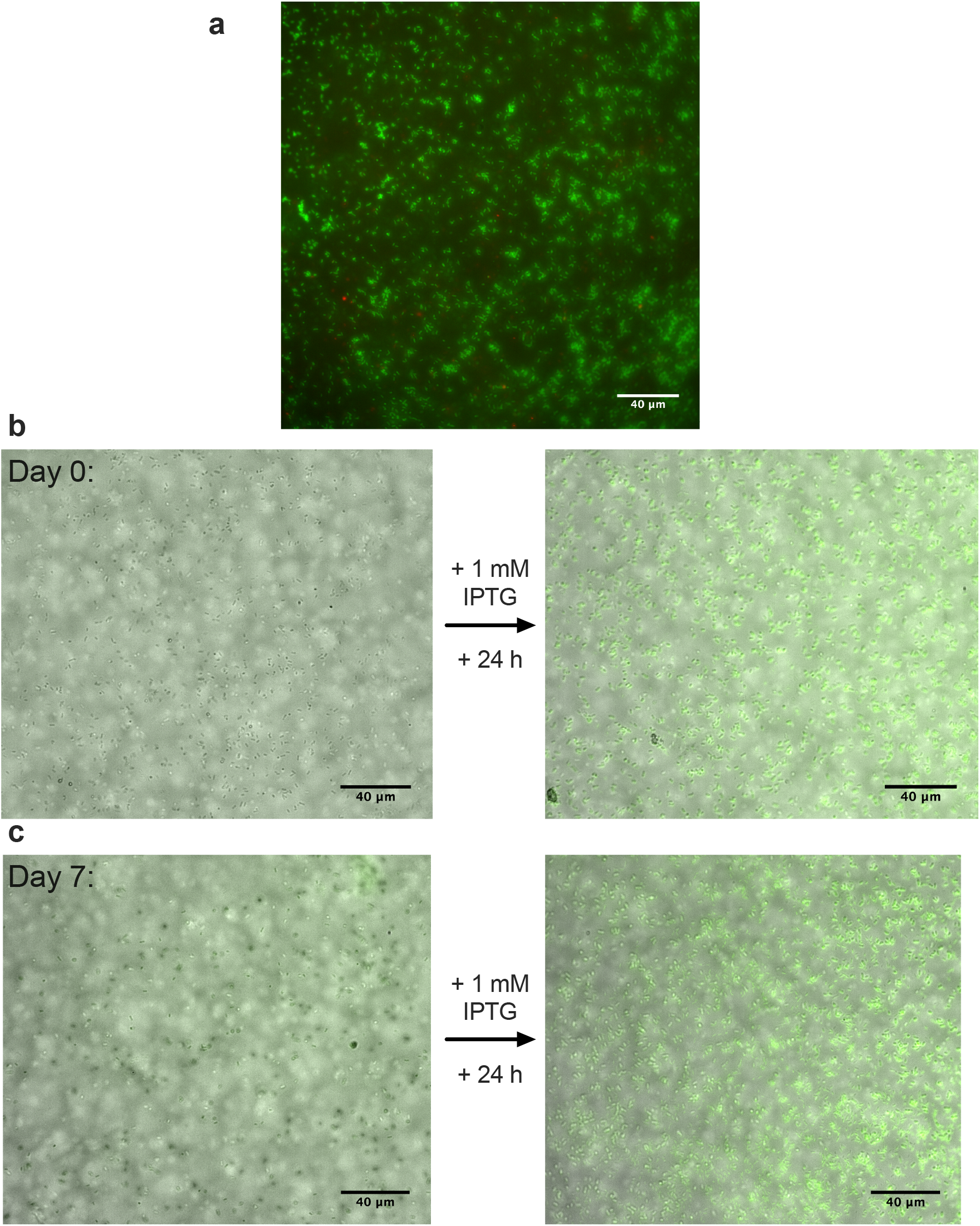
*S. oneidensis* retains viability after cross-linking and remains metabolically active within hydrogels for at least one week. **a**, BacLight live/dead staining of *S. oneidensis* MR-1 after cross-linking, swelling overnight in 1x PBS, and mechanical characterization by rheology. Cells are predominantly alive (green fluorescence) as opposed to dead (red fluorescence). **b–c**, Overlaid fluorescence and bright-field microscopy of *S. oneidensis* MR-1 + *sfgfp* (left) encased in gels **b**, one day or **c**, one week after cross-linking and swelling, and (right) 24 h after inoculating with 1 mM IPTG to induce fluorescence. Images are representative of *n* = 3 biological replicates.

**Extended Data Fig. 3.**
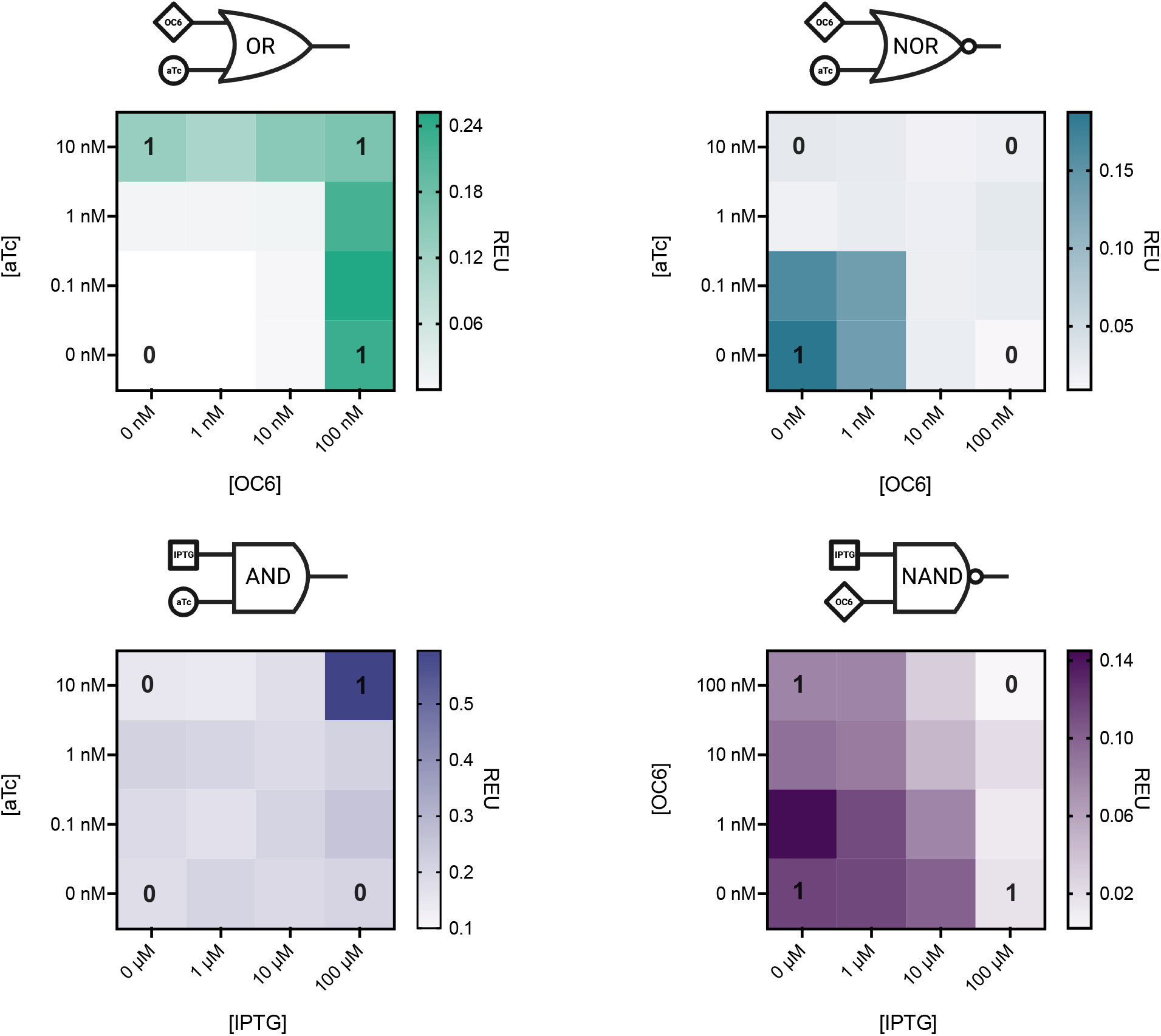
Genetic Boolean logic enables concentration-dependent transcriptional responses in *S. oneidensis* when expressing *sfgfp*. Relative Expression Units (REU) measured as a function of combinatorial inducer concentration show characteristic transcriptional regulation that follows expected truth tables for multiple genetic Boolean architectures expressing *sfgfp* (OR, NOR, NAND) or *eYFP* (AND). *S. oneidensis* MR-1 harboring each plasmid was grown overnight in aerobically prepared 96-well plates that were then sealed to emulate dynamic cross-linking conditions. Fluorescence was OD_600_-normalized and referenced to a constitutive fluorescence plasmid to obtain REU. The expected truth tables are shown for maximum and minimum induction conditions (“0”, OFF; “1”, ON). Data shown are mean of *n* = 3 biological replicates.

**Extended Data Fig. 4.**
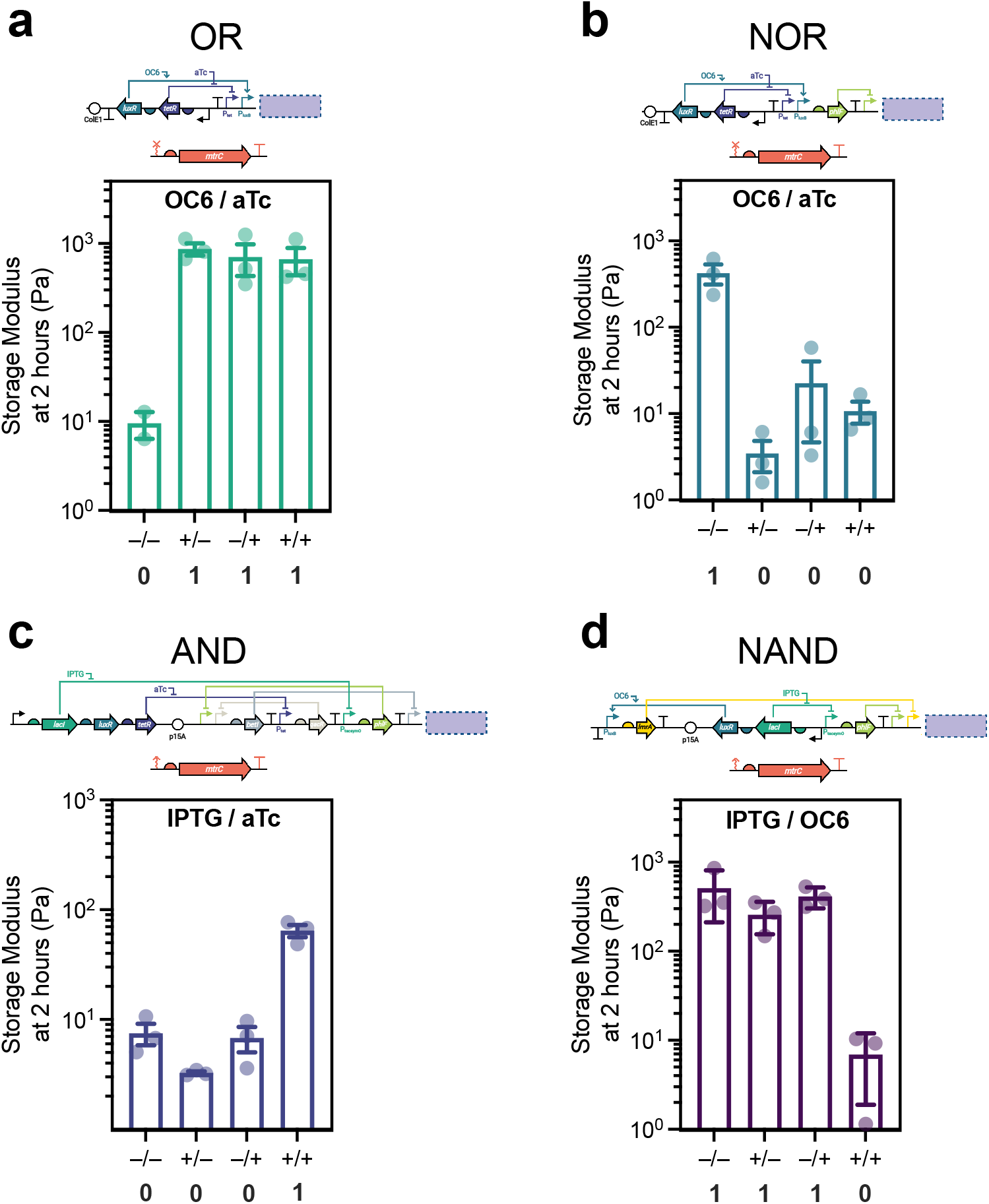
Genetic Boolean logic enables synthetic material computation during stationary phase cross-linking. Storage moduli for networks cross-linked using *S. oneidensis* strains harboring transcriptional Boolean logic circuits controlling *mtrC* expression. The logical architectures span **a**, OR, **b**, NOR, **c**, AND, and **d**, NAND using nested repressors that coordinate in response to sensor blocks. The repressors are activated or deactivated in response to their cognate inducing molecules (100 μM IPTG, 100 nM OC6, 10 nM aTc, and/or their corresponding solvent blanks). In all cases, storage modulus was measured 2 h after inoculation. The expected truth tables are represented below each circuit (“0”, OFF; “1”, ON). Each network appropriately responds to combinatorial inputs by increasing/decreasing storage modulus in response to *sfgfp* or *mtrC* activation/deactivation. Each plasmid architecture is shown as a cartoon above the corresponding response function, with representations as in Figure 1. Data shown are mean ± SEM of *n* = 3 biological replicates.

**Extended Data Fig. 5.**
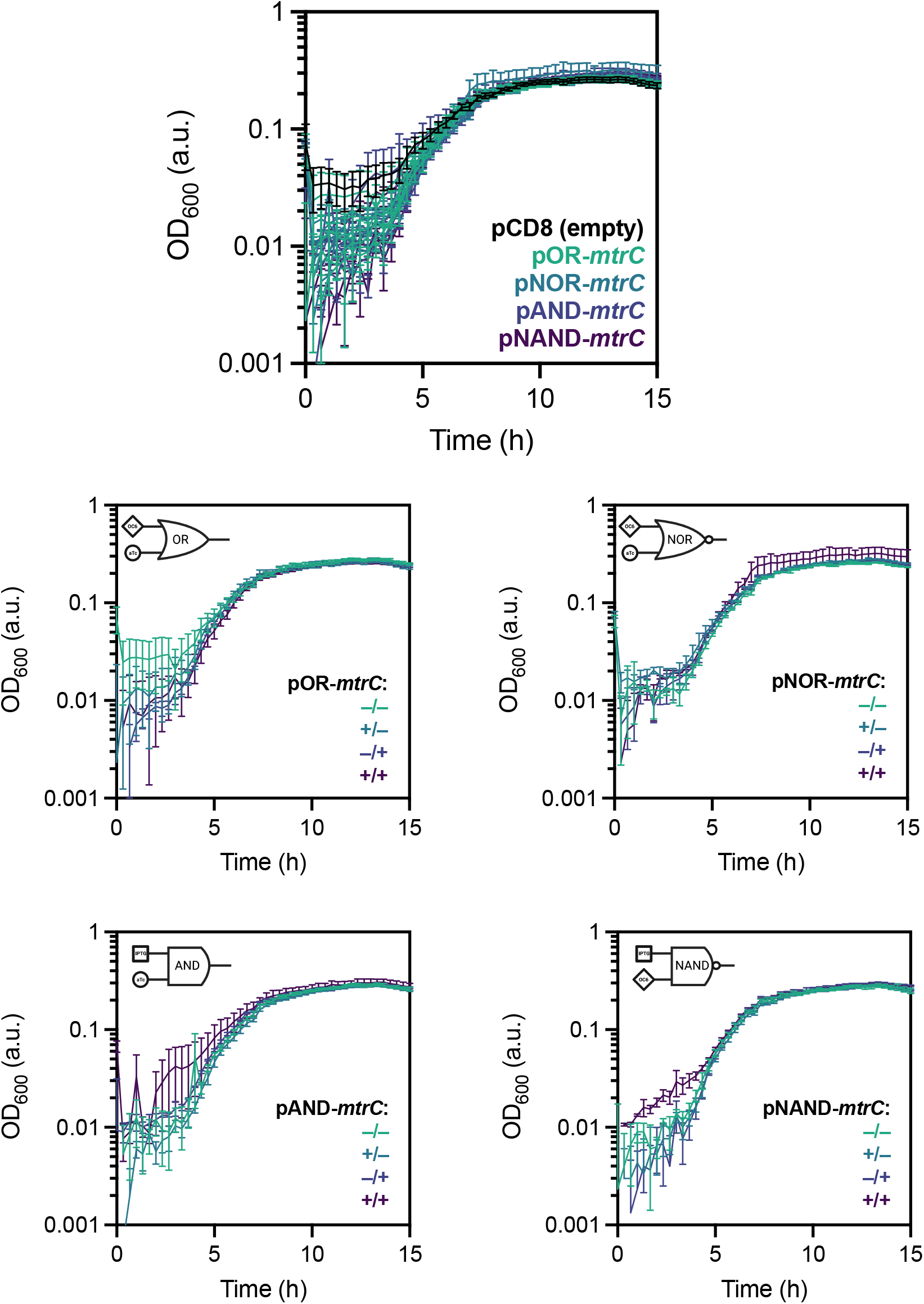
Growth kinetics are not affected by induction in *S. oneidensis* harboring genetic Boolean logic controlling *mtrC*. OD_600_ measured *in situ* at 30 °C for *S. oneidensis* harboring each Boolean *mtrC* construct under varying inducer conditions. An induced empty vector control (pCD8) was also measured as a reference. In general, growth was not affected by induction. Data shown are mean ± SEM of *n* = 3 biological replicates.

**Extended Data Fig. 6.**
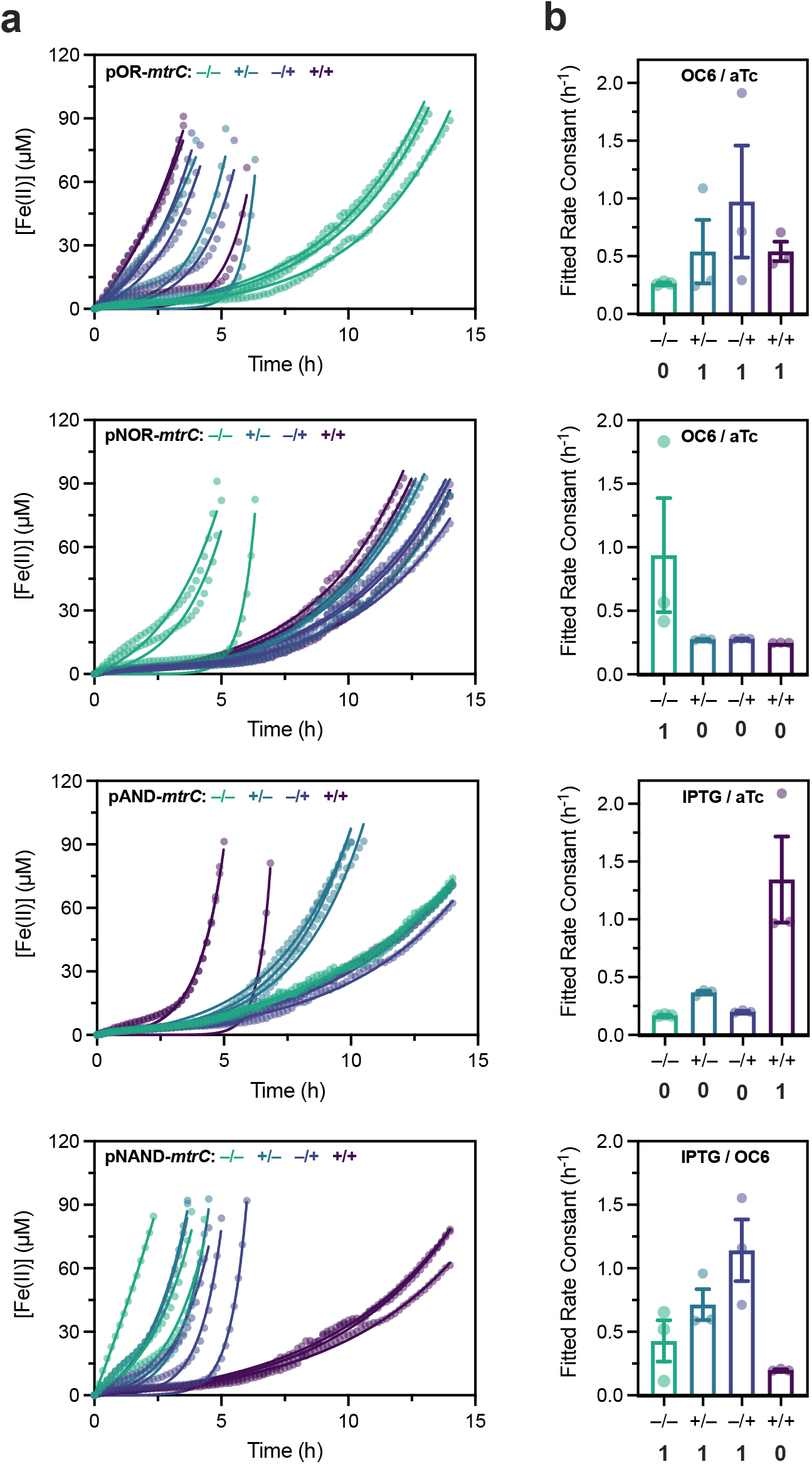
Genetic Boolean logic enables input signal-dependent metal reduction. **a**, Raw *in situ* Fe(III) reduction kinetics measured using ferrozine absorbance and corresponding Monod-type fits^40^. **b**, Fitted Fe(III) reduction rate constants for corresponding curves calculated using a Monod-type model, with expected truth tables shown below (“0”, OFF; “1”, ON). Data shown are mean ± SEM of *n* = 3 biological replicates.

**Extended Data Fig. 7.**
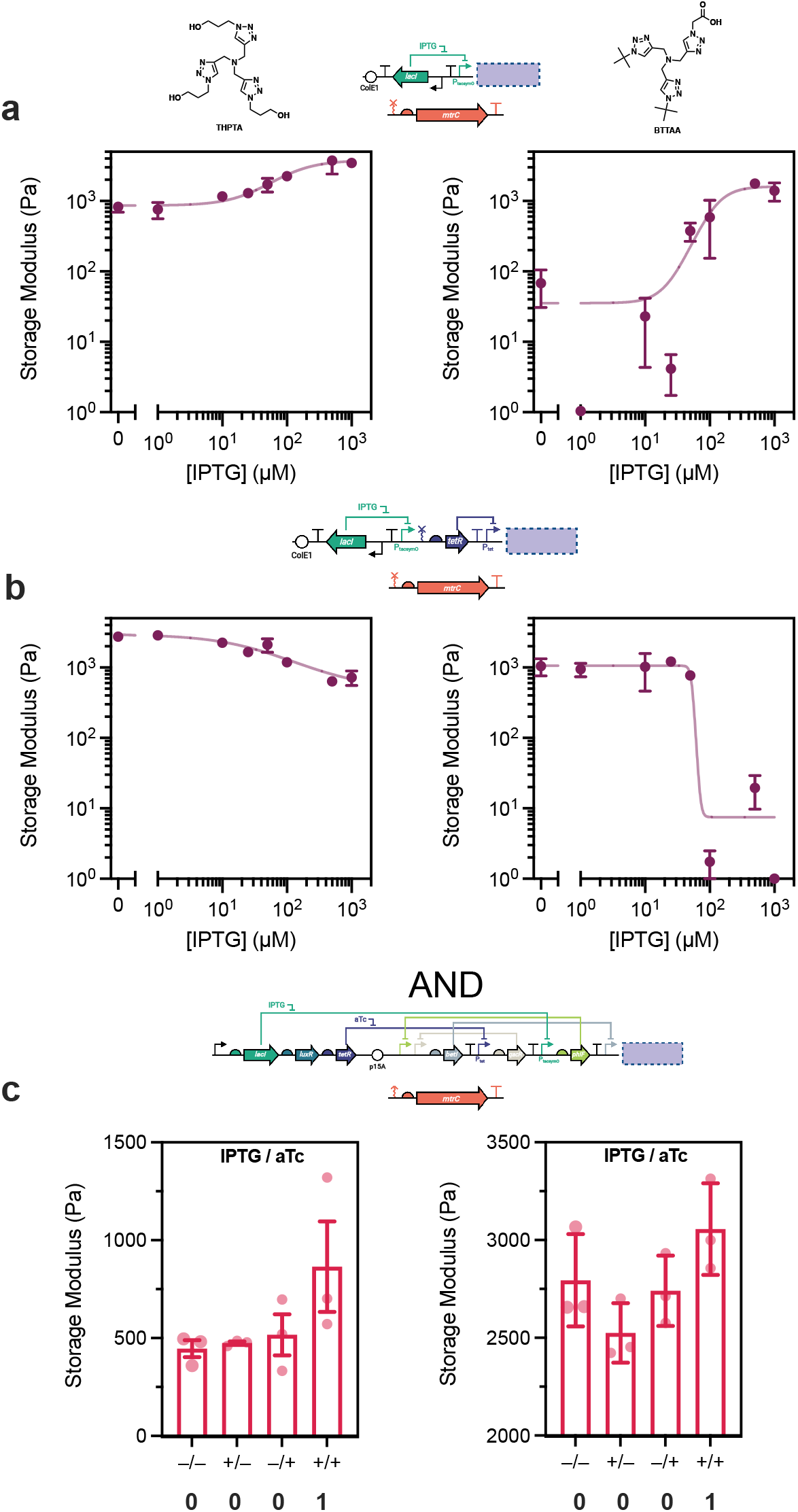
Dynamic range and ON/OFF behavior of EET-controlled CuAAC is modulated by the Cu ligand. Storage moduli and corresponding response functions for networks cross-linking using *S. oneidensis* harboring **a**, Buffer, **b**, NOT, and **c**, AND gate architectures controlling *mtrC* expression, with either THPTA (left) or BTTAA (right) as the Cu ligand. Storage modulus was measured 16 h (THPTA) or 12 h (BTTAA) after inoculation. The expected truth tables are represented below each AND circuit (“0”, OFF; “1”, ON). Chemical structures of each ligand are represented above; each plasmid architecture is shown as a cartoon above the corresponding response function, with representations as in Figure 1. THPTA, tris(benzyltriazolylmethyl)amine; BTTAA, 2-(4-((bis((1-(tert-butyl)-1H-1,2,3-triazol-4-yl)methyl)amino)methyl)-1H-1,2,3-triazol-1-yl)acetic acid.

