## Supplementary Information for "Transcriptional Regulation of Synthetic Polymer Networks"

#### Methods

##### *Chemicals and Reagents*

4-arm 5k poly(ethylene glycol) methacrylate (PEG-MA, ≥95% functionalization, Advanced BioChemicals), copper(II) bromide ($CuBr_2$, Sigma-Aldrich, 99%), tris(2-pyridylmethyl)amine (TPMA, Sigma-Aldrich, 98%), Tris(benzyltriazolylmethyl)amine (THPTA, Sigma-Aldrich, 95%), 2-(4-((bis((1-(tert-butyl)-1H-1,2,3-triazol-4-yl)methyl)amino)methyl)-1H-1,2,3-triazol-1-yl)acetic acid (BTTAA, Click Chemistry Tools > 95%), 2-hydroxyethyl 2-bromoisobutyrate (HEBIB, Sigma-Aldrich, 95%), sodium DL-lactate ($NaC_3H_5O_3$, TCI, 60% in water), sodium fumarate ($Na_2C_4H_2O_4$, VWR, 98%), HEPES buffer solution ($C_8H_{18}N2O_4S$, VWR, 1 M in water, pH = 7.3), potassium phosphate dibasic ($K_2HPO_4$, Sigma-Aldrich), potassium phosphate monobasic ($KH_2PO_4$, Sigma-Aldrich), sodium chloride (NaCl, VWR), ammonium sulfate ($(NH_4)_2SO4$, Fisher Scientific), magnesium(II) sulfate heptahydrate ($MgSO_4 \cdot 7H2O$, VWR), trace mineral supplement (ATCC), casamino acids (VWR), isopropyl ß-D-1-thiogalactopyranoside (IPTG, Teknova), anhydrotetracycline hydrochloride (aTc, Sigma-Aldrich), 3-oxohexanoyl-homoserine lactone (OC6, Sigma-Aldrich), kanamycin sulfate ($C_{18}H_{38}N_4O_{15}S$, Growcells), iron(III) citrate ($C_6H_5FeO_7$, Alfa Aesar), iron (II) sulfate heptahydrate ($FeSO_4 \cdot 7H_2O$, Alfa Aesar), 3-(2-Pyridyl)-5,6-bis(4-sulfophenyl)-1,2,4-triazine disodium salt hydrate (ferrozine, $C_{20}H_{12}N_4Na_2O_6S_2$, TCI), 5-norbornene-2-carboxylic acid (Sigma-Aldrich), N,N'-diisopropylcarbodiimide (Sigma-Aldrich), 4-dimethylaminopyridine (Sigma-Aldrich), diethyl ether (Acros Organics), anhydrous DCM, DMSO, and $CDCl_3$ (Sigma-Aldrich), DMSO-$d^6$ (Sigma-Aldrich), nail polish (Electron Microscopy Sciences), BacLight Live/Dead Stain (Invitrogen), deuterium oxide ($D_2O$, Sigma-Aldrich, 99.9%), were used as received. All other poly (ethylene-glycol) macromers were purchased from Jenkem USA. All media components were autoclaved or sterilized using 0.2 µm PES filters.

##### *Bacteria Strains and Culture*

Bacterial strains and plasmids are listed in Table S1. Cultures were prepared as follows: bacterial stocks stored in 20% glycerol at −80 °C were streaked onto LB agar plates (for wild-type and knockout strains) or LB agar with 20 or 25 µg/mL kanamycin (for plasmid-harboring strains) and grown overnight at 30 °C for *Shewanella* and 37 °C for *E. coli*. Single colonies were isolated and inoculated into *Shewanella* Basal Medium (SBM) supplemented with 100 mM HEPES, 0.05% trace mineral supplement, 0.05% casamino acids, and 20 mM sodium lactate (2.85 µL of 60% w/w sodium lactate per 1 mL culture) as the electron donor. Aerobic cultures were pregrown in 15 mL culture tubes at 30 °C and 250 rpm shaking. Anaerobic cultures were pregrown using the same procedure outlined above, but in degassed growth medium in a humidified anaerobic chamber (3% $H_2$, balance $N_2$, Coy) and supplemented with 40 mM sodium fumarate (40 µL of 1 M sodium fumarate per 1 mL culture) as the electron acceptor. For stationary phase conditions, inducible strains were pregrown anaerobically without inducer(s) for 4–6 h before being diluted 1:25 into inducer-containing media (from 1000x stocks) to grow overnight. Cultures were washed 3x after pregrowth using SBM supplemented with 0.05% casamino acids (degassed for anaerobic cultures). $OD_{600}$ was measured using a NanoDrop 2000C spectrophotometer and normalized to 10x the inoculating $OD_{600}$ for dilution into gel mixtures (5 µL of concentrated cell culture into 45 µL of gel mixture) unless otherwise noted.

##### *Hydrogel Radical Cross-Linking Procedure using Engineered S. oneidensis*

$CuBr_2$ and TPMA were dissolved at 8 mM in DMF for storage and combined into a 400 µM Cu-TPMA stock solution in DMF before reaction preparation. HEBIB (1.45 µL) was added to SBM with casamino acids (143 µL) to create a 69 mM stock solution which was diluted 5-fold in SBM with casamino acids to create a 13.8 mM solution before reaction preparation. Per 50 µL hydrogel disc that was analyzed by rheology, a cross-linking reaction mixture was prepared as follows: PEG-MA was dissolved at 6.18 wt % in SBM with 0.05% casamino acids and aliquoted into an autoclaved microfuge tube (40.47 µL). Solutions of 400 µM Cu-TPMA (0.625 µL or 1.25 µL), 13.8 mM HEBIB (0.3625 µL), 60% sodium lactate (0.143 µL), and 1 M sodium fumarate (2 µL) were added to the PEG-MA solution and mixed. Per 50 µL gel mixture, the remaining 1.4 µL was used for antibiotic and inducing molecule addition where necessary, or to compensate for varying Cu-TPMA concentration, otherwise 1.4 µL of SBM with casamino acids was added. Constituent volumes were multiplied as necessary to create a single primary stock for each experiment involving identical inducer conditions and *S. oneidensis* strains. The final concentrations in solution were 5 wt % PEG-MA, 5 or 10 µM Cu-TPMA, 100 µM HEBIB, 20 mM lactate, 40 mM fumarate, and 0, 20, or 25 µg/mL kanamycin where necessary, depending on *S. oneidensis* strain. Inducer concentrations ranged from 0 to 1000 µM depending on the condition and were diluted from 100x stock solutions. IPTG was dissolved in sterile $H_2O$, aTc was dissolved in a 1:1 ethanol:$H_2O$ solution, and OC6 was dissolved in DMF; all inducer stocks were stored at −20 °C. The primary gel mixture was then distributed into individual autoclaved microfuge tubes of 45 µL aliquots to which 5 µL of $OD_{600}$-normalized cells were added. The gel solutions were mixed and dispensed onto hydrophobically treated glass slides with a 0.5 mm silicone spacer separating the two glass layers. The gels were allowed to react at 30 °C for 2 h at inoculating $OD_{600}$ = 0.2 (stationary phase conditions), or 16 to 24 h at inoculating $OD_{600}$ values ranging from 0.01 to 0.05, depending on the strain (dynamic conditions). Hydrogels were removed from the slides using a razor blade and placed into 3 mL baths of 1x PBS overnight to swell to equilibrium at room temperature in the dark.

##### *Rheological Analysis*

Swollen hydrogels prepared as outlined above were analyzed by oscillatory shear rheology using a TA Instruments Discovery HR-2 Rheometer with an 8 mm parallel plate geometry. Hydrogels were loaded onto a Peltier plate and excised to 8 mm diameter using a biopsy punch. The geometry gap was then lowered until the measured axial force remained at or above 0.02 N where possible (usually between 300−600 µm, depending on the crosslink density and swelling ratio). Storage and loss moduli were measured using frequency sweeps from 0.1 to 1 Hz at a constant strain of 1%. Moduli for a single gel were quantified by averaging the linear viscoelastic region of each frequency sweep.

##### *Plasmid Construction*

All bacterial strains, plasmids, genetic circuit maps, and sequence information for each genetic part are detailed in Tables S1–2, 4. All plasmids were purchased from Addgene or assembled via Golden Gate cloning procedures using enzymes (BsaI, SapI, BsmBI) and buffers from New England Biolabs. DNA fragments used in Golden Gate cloning were generated via partial/whole-plasmid PCR or commercially synthesized (Integrated DNA Technologies or Twist Biosciences). Generally, 10 µL Golden Gate reactions were set up that contained 10 fmol of plasmid backbone and 40 fmol of each synthesized gene and/or PCR insert (as necessary). In a thermocycler, Golden Gate reactions were cycled 25-45 times, depending on the complexity and size of the construct: 90 s at 37 °C (for BsaI and SapI) or 42 °C (for BsmBI) followed by 3 min at 16 °C. After the cycles, reactions were incubated at 37 °C (for BsaI and SapI) or 55 °C (for BsmBI) for 5 min, 80 °C for 10 min, and then held at 4 °C. Golden Gate reactions were used to directly transform freshly prepared electrocompetent *S. oneidensis* or *E. coli* strains. To prepare electrocompetent *S. oneidensis*, 5 mL of overnight *S. oneidensis* growth in LB medium at 30 °C was washed 3 times with sterile 10% glycerol at room temperature and concentrated to ∼300 µL. A 2 µL portion of Golden Gate reaction was mixed with 30 µL of concentrated electrocompetent *S. oneidensis*, transferred to a 1 mm electroporation cuvette, and electroporated at 1250 V. To recover electroporated cells, 250 µL of LB warmed in a 30 °C incubator was immediately added post-electroporation and cells were incubated/shaken at 30 °C and 250 rpm. After 2 h of recovery, 100 µL of cell suspension was plated onto LB agar plates containing 20 or 25 µg • $mL^{-1}$ kanamycin and incubated overnight at 30 °C to obtain single colonies. Single colonies were used to inoculate LB liquid medium containing 20 or 25 µg • $mL^{-1}$ kanamycin sulfate and incubated/shaken overnight at 30 °C and 250 rpm. These cultures were used to generate 20–22.5% glycerol stocks which were stored at −80 °C, and to harvest assembled plasmid for Sanger sequencing (DNA Sequencing Facilities, University of Texas at Austin).

##### *Functional Verification of* mtrC *Expression*

Strains containing *mtrC* expression circuits were functionally validated using an *in situ* Fe(III) reduction/ferrozine assay as previously described[1]. Briefly, strains were anaerobically pregrown for ca. 6 h in SBM containing 20 mM lactate, 40 mM fumarate, and 20 or 25 µg • $mL^{-1}$ kanamycin depending on the strain. These cell suspensions were diluted 100-fold into SBM containing 20 mM lactate, 40 mM fumarate, kanamycin, and appropriate inducers, then allowed to grow for ca. 18 h. Subsequently, these growths were diluted 100-fold into 96-well plates containing SBM solution with 20 mM lactate, kanamycin, 1 mg • $mL^{-1}$ ferrozine, appropriate inducers, and 5 mM Fe(III) citrate, such that the final well volume was 250 µL. Fe(II) standards were also included in the plate using dissolved $FeSO_4$. The 96-well plate was sealed with a sterile/optically transparent film (PCR-SP-S, AxySeal Scientific), covered with a polystyrene plate lid (Eppendorf) with silicone grease lining the edges, removed from the anaerobic chamber, and placed within a BMG LABTECH CLARIOstar plate reader with temperature control set to 30 °C. Without shaking, the absorbance at 562 nm was measured every 10 min for at least 14 h. Using the Fe(II) standards, raw kinetics data was converted to Fe(II) concentrations vs. time. Fe(II) kinetics for individual replicates were background subtracted (i.e. Fe(II) level at the initial time point) and fitted to an exponential Monod-type model to obtain fitted rate constants ($\mu$): $\mathrm{Fe}_{\mathrm{subt}}^{\mathrm{II}} = K(\exp(\mu t) - 1$.

##### *Quantification and Modeling of Fluorescence and Cross-Linking Constructs*

Strains containing *sfgfp* architectures were aerobically pregrown overnight using standard culture conditions outlined above, with the addition of 20 or 25 µg • $mL^{-1}$ kanamycin depending on the strain. Cultures were then diluted 1:25 into 96-well plates containing SBM with 0.05% casamino acids, 20 mM lactate, 40 mM fumarate, kanamycin, 5 µM Cu-TPMA, 100 µM HEBIB, and varying amounts of inducer(s) (from 500x stocks). Plates were sealed with impermeable foil and placed at 30 °C for 18–24 h. Prior to measuring sfGFP fluorescence, protein translation was arrested by supplementing a 100 µL aliquot of cell suspension with kanamycin sulfate to a final concentration of 2 mg • $mL^{-1}$. Subsequently, this suspension was shaken aerobically for 1 h at 30 °C to allow for sfGFP maturation. sfGFP fluorescence (488/530 nm) and cell suspension absorbance (600 nm) were measured using a BMG LABTECH CLARIOstar plate reader to yield fluorescence • $absorbance^{-1}$ for each sample. For each sample, the background fluorescence • $absorbance^{-1}$ from an empty vector (pCD8) control was subtracted. In addition, strains were normalized in each plate to a RNAP flux standard strain constitutively expressing *sfgfp* (pCDe1) via the Ptrc* promoter to enable relative expression unit (REU) calculations. A nonlinear fitting algorithm in GraphPad Prism 9 was used to fit inducible gene expression to the following activating Hill function: $y = \min + (\max - \min)\frac{[I]^n}{{K_{1/2}}^n + [I]^n}$. Strains containing *mtrC* architectures were analyzed similarly after rheological analysis to fit hydrogel storage modulus. Normalized hydrogel storage modulus was calculated using the average storage modulus from gels cross-linked using wild-type *S. oneidensis* harboring a representative empty vector plasmid (pCD8). Further details on modeling can be found in previous work[1,2]. Fitting parameters and "goodness of fit" can be found in Table S3.

##### *Kinetic Growth Measurements*

Genetic Boolean logic strains containing *mtrC* architectures were aerobically pregrown overnight using standard culture conditions outlined above, with the addition of 20 µg • $mL^{-1}$ kanamycin. Cultures were then diluted 1:100 into 96-well plates containing SBM with 0.05% casamino acids, 20 mM lactate, 40 mM fumarate, kanamycin, and varying combinations of appropriate inducer(s) and blanks (from 500x stocks) for a total volume of 250 µL. Plates were sealed with impermeable foil and placed at 30 °C for ca. 18 h, and $OD_{600}$ was tracked using a BMG LABTECH CLARIOstar plate reader. An empty vector control (pCD8) induced with 1000 µM IPTG was also measured as a reference.

##### *CuAAC Hydrogel Cross-Linking*

$CuBr_2$ was dissolved at 8 mM in DMF for storage and combined into with THPTA or BTTAA in sterile water to form a 500 µM Cu-ligand stock solution a ratio of 1:6 Cu:ligand. Per 50 µL hydrogel disc that was analyzed by rheology, a cross-linking reaction mixture was prepared as follows: 4-arm-PEG-alkyne (5K) was dissolved at 4.62 wt % in SBM with 0.05% casamino acids and aliquoted into an autoclaved microfuge tube (18.68 µL) where it was combined with 4-arm-PEG-azide (10K) was dissolved at 8.92 wt % in SBM with 0.05% casamino acids (18.68 µL). Solutions of 500 µM Cu-THPTA (5 µL), 60% sodium lactate (0.143 µL), and 1 M sodium fumarate (1 µL) were added to the solution and mixed. Per 50 µL gel mixture, the remaining 1.5 µL was used for antibiotic and inducing molecule addition where necessary, otherwise 1.5 µL of SBM with casamino acids was added. Constituent volumes were multiplied as necessary to create a single primary stock for each experiment involving identical inducer conditions and *S. oneidensis* strains. The final concentrations in solution were 5 wt % PEG-backbone (1.67 wt % 4-arm-PEG-alkyne, and 3.33 wt % 4-arm-PEG-azide), 50 µM Cu-THPTA (1:6) or Cu-BTTAA (1:6), 20 mM lactate, 20 mM fumarate, and 25 µg/mL kanamycin where necessary, depending on *S. oneidensis* strain. Inducer concentrations ranged from 0 to 1000 µM depending on the condition and were diluted from 100x or 200x stock solutions. The primary gel mixture was then distributed into individual autoclaved microfuge tubes of 45 µL aliquots to which 5 µL of $OD_{600}$-normalized cells were added. The gel solutions were mixed and dispensed onto hydrophobically treated glass slides with a 0.5 mm silicone spacer separating the two glass layers. The gels were allowed to react at 30 °C for 12 to 16 h at inoculating $OD_{600}$ values ranging from 0.01 to 0.05, depending on the strain (dynamic conditions). Hydrogels were removed from the slides using a razor blade and placed into 3 mL baths of 1x PBS overnight to swell to equilibrium at room temperature.

##### *Synthesis of Poly(ethylene-glycol) Alkyne (PEG-Alk)*

PEG-Alk was synthesized by first adding 0.2 mmol of 4 arm, 5 kDa PEG-Acetic Acid and 2.8 mmol of HBTU to a 25 mL oven dried round bottom flask. 5 mL of anhydrous DMSO was added. 0.88 mmol of propargylamine and 2.8 mmol of triethylamine were added. The flask was sealed with a rubber septa and purged with argon for 5 minutes. The reaction was allowed to proceed for 24 hours. Then, the solution was precipitated into 35 mL of ice-cold diethyl ether. The resulting ether was decanted, and the white powder was dissolved in pure water and placed into a 1 kDa molecular weight cut-off dialysis bag. The product was dialyzed in 4L of pure water for 3 days, changing the water once a day. After 3 days, the product dissolved in water was lyophilized to yield a white powder product. Yield=99%. $^1H$ NMR ($CDCl_3$, 400 MHz): δ=2.22 (t, 1H), δ=3.62 (m, 113H), δ=4.00 (s, 2H), δ=4.08 (dd, 2H), δ=7.42 (s, 1H). NMR shows approximately a 90% functionalization.

##### *Microscopy*

Microscopy was performed using a Nikon Ti2 Eclipse inverted epifluorescence microscope. Cells assessed for viability by microscopy were cross-linked using standard conditions and the resulting gels swollen in 1x PBS at room temperature overnight. The gels were then incubated in the dark in a BacLight Live/Dead stain mix (1.5 µL/mL Syto9, 2.5 µL/mL propidium iodide in 0.85% NaCl solution) for 30 min. Stained gels were then washed by pipetting 3x in 1 mL PBS to remove unbound dye. Gels were loaded onto glass microscope slides, and a no. 1 coverslip was placed on top. The gel thickness prevented using nail polish to seal the sides, but evaporative losses were not noticeable over the course of the experiment (~30 min).

Fluorescence for each stain (green for Syto9, red for propidium iodide) was measured using GFP and Texas Red excitation/emission filter cubes on a Nikon Ti2 Eclipse. To assess metabolic activity, gels were cross-linked with *sfgfp*-harboring strains and allowed to swell in 1x PBS. sfGFP fluorescence was assessed before induction to measure background fluorescence. Gels were then incubated in 0 μM or 1000 μM IPTG in PBS for 24 h and monitored by fluorescence using the GFP channel.

**Table S1.** Bacterial strains and plasmids used in this study.

| Strain or plasmid | Description/Genotype | Reference or source |
|---|---|---|
| ***S. oneidensis* Strains (+plasmid)** | | |
| MR-1 | MR-1 (ATCC700550), wild-type strain | American-Type Culture Collection |
| JG596 | Lacks outer membrane cytochromes MtrC, OmcA, and MtrF; Δ*mtrC*Δ*omcA*Δ*mtrF* | Jeffrey Gralnick, U. of Minnesota |
| Δ*cymA* | Lacks inner membrane cytochrome CymA | Jeffrey Gralnick, U. of Minnesota |
| Δ*mtrA* | Lacks periplasmic cytochrome, MtrA | [1] |
| MR-1+pCDe1 | Wild-type with *sfgfp* REU plasmid | This work |
| MR-1+pCDhCK7 | Wild-type with *sfgfp* REU plasmid | This work |
| MR-1+pCD7sfGFP | Wild-type with *sfgfp* Buffer gate | [1] |
| MR-1+pCD8 | Wild-type with empty Buffer gate | [1] |
| MR-1+pCDhCK8 | Wild-type with empty MoClo backbone | This work |
| Δ*cymA*+pCD8 | Δ*cymA* with empty Buffer gate | [1] |
| Δ*cymA*+pCD26r4 | Δ*cymA* with *cymA* Buffer gate (sRBS4$_{cymA}$) | [1] |
| Δ*mtrA*+pCD8 | Δ*mtrA* with empty Buffer gate | [1] |
| Δ*mtrA*+pCD25r0 | Δ*mtrA* with *mtrA* Buffer gate (sRBS0$_{mtrA}$) | [1] |
| JG596+pCD8 | JG596 with empty Buffer gate | [1] |
| JG596+pCD24r1 | JG596 with *mtrC* Buffer gate (sRBS1$_{mtrC}$) | [1] |
| JG596+pCDd1 | JG596 with *mtrC* NOT gate (sRBS1$_{mtrC}$) | [1] |
| MR-1+pCDTetRsfGFP | Wild-type with *TetR*-regulated *sfgfp* | This work |
| JG596+pCD24tr1 | JG596 with *TetR*-regulated *mtrC* | This work |
| MR-1+pAT3 | Wild-type with *LuxR*-regulated *sfgfp* | This work |
| JG596+pAT1 | JG596 with *LuxR*-regulated *mtrC* | This work |
| MR-1+pIEM12-*sfgfp* | Wild-type with T7-regulated *sfgfp* | This work |
| JG596+pIEM12 | Wild-type with T7-regulated *mtrC* | This work |
| MR-1+pCDhCK4 | Wild-type with IPTG-regulated *sfgfp* anti-repressor NOT gate | This work |
| JG596+pCDhCK3 | JG596 with IPTG-regulated *mtrC* anti-repressor NOT gate | This work |
| MR-1+pCDhCK6 | Wild-type with ribose-regulated *sfgfp* anti-repressor NOT gate | This work |
| JG596+pCDhCK5 | JG596 with ribose-regulated *mtrC* anti-repressor NOT gate | This work |
| MR-1+pAJGb1-*sfgfp* | Wild-type with *sfgfp* OR gate | This work |
| JG596+pAJGb1-*mtrC* | JG596 with *mtrC* OR gate | This work |
| MR-1+pAJGb2-*sfgfp* | Wild-type with *sfgfp* NOR gate | This work |
| JG596+pAJGb2-*mtrC* | JG596 with *mtrC* NOR gate | This work |
| MR-1+pAJGb3-*sfgfp* | Wild-type with *sfgfp* AND gate | This work |
| JG596+pAJGb3-*mtrC* | JG596 with *mtrC* AND gate | This work |
| MR-1+pAJGb4-*sfgfp* | Wild-type with *sfgfp* NAND gate | This work |
| JG596+pAJGb4-*mtrC* | JG596 with *mtrC* NAND gate | This work |
| **Plasmids** | | |
| pCDe1 | *sfgfp* constitutive REU (Ptrc* promoter) | This work |

| | | |
|---|---|---|
| pCD7sfGFP | *sfgfp* Buffer gate ($P_{tacsym0}$; *lacI*) | [1] |
| pCD8 | empty Buffer gate ($P_{tacsym0}$; *lacI*) | [1] |
| pCD24r1 | *mtrC* Buffer gate ($P_{tacsym0}$; *lacI*) | [1] |
| pCD25r0 | *mtrA* Buffer gate ($P_{tacsym0}$; *lacI*) | [1] |
| pCD26r4 | *cymA* Buffer gate ($P_{tacsym0}$; *lacI*) | [1] |
| pCDd1 | *mtC* NOT gate ($P_{tacsym0}$; *lacI*) | [1] |
| pCD24tr1 | *mtrC* Buffer gate ($P_{Tet}$; *tetR*) | This work |
| pCDTetRsfGFP | *sfgfp* Buffer gate ($P_{Tet}$; *tetR*) | This work |
| pAT1 | *mtrC* Buffer gate ($P_{Lux}$; *luxR*) | This work |
| pAT3 | *sfgfp* Buffer gate ($P_{Lux}$; *luxR*) | This work |
| pAJGb1-*mtrC* | *mtrC* OR gate (($P_{Tet}$; *tetR*; $P_{Lux}$; *luxR*) | This work |
| pAJGb1-*sfgfp* | *sfgfp* OR gate (($P_{Tet}$; *tetR*; $P_{Lux}$; *luxR*) | This work |
| pAJGb2-*mtrC* | *mtrC* NOR gate (($P_{Tet}$; *tetR*; $P_{Lux}$; *luxR*) | This work |
| pAJGb2-*sfgfp* | *sfgfp* NOR gate (($P_{Tet}$; *tetR*; $P_{Lux}$; *luxR*) | This work |
| pAJGb3-*mtrC* | *mtrC* AND gate ($P_{tacsym0}$; *lacI*; $P_{Tet}$; *tetR*) | This work; [3] |
| pAJGb3-*eYFP* | *eYFP* AND gate ($P_{tacsym0}$; *lacI*; $P_{Tet}$; *tetR*) | This work; [3] |
| pAJGb4-*mtrC* | *mtrC* NAND gate ($P_{tacsym0}$; *lacI*; $P_{Lux}$; *luxR*) | This work; [3] |
| pAJGb4-*sfgfp* | *sfgfp* NAND gate ($P_{tacsym0}$; *lacI*; $P_{Lux}$; *luxR*) | This work; [3] |

**Table S2.** Genetic parts/sequences used to construct the plasmids in this study.

Underline indicates added insulator sequence.

| Genetic Part | DNA Sequence (5' to 3') |
|---|---|
| **Promoters** | |
| $P_{tacsymO}$[4] | TGTTGACAATTAATCATCGGCTCGTATAATGTGTGGAATTGTGAGCGCTCACAATTCTATGGACTATGTTT |
| $P_{lacI}$ | GCGGCGCGCCATCGAATGGCGCAAAACCTTTCGCGGTATGGCATGATAGCGCCCGGAAGAGAGTCAATTCAGGGTGGTGAAT |
| $P_{lacIQ}$[5] | GCGGCGCGCCATCGAATGGTGCAAAACCTTTCGCGGTATGGCATGATAGCGCCC |
| $P_{tet}$[6] | TACTCCACCGTTGGCTTTTTTCCCTATCAGTGATAGAGATTGACATCCCTATCAGTGATAGAGATAATGAGCAC |
| $P_{luxB}$ | GCTTAACGATCGTTGGCTGACCTGTAGGATCGTACAGGTTTACGCAAGAAAATGGTTTGTTACAGTCGAATAAA |
| $P_{trc*}$[7] | GAAATATTCTGAAATGAGCTGTTGACAATTAATCATCCGGCTCGTATAATGTGTGGAA |
| $P_{phlF}$[3] | CGACGTACGGTGGAATCTGATTCGTTACCAATTGACATGATACGAAACGTACCGTATCGTTAAGGT |
| $P_{qacR}$[3] | GGTATGGAAGCTATACGTTACCAATTGACAGCTAGCTCAGTCCTACTTTAGTATATAGACCGTGCGATCGGTCTATA |
| $P_{betI}$ [3] | AGCGCGGGTGAGAGGGATTCGTTACCAATTGACAATTGATTGGACGTTCAATATAATGCTAGC |
| $P_{lmrA}$ [3] | CGCTCATTCACTAGGTCTGATTCGTTACCAATTGACAACTGGTGGTCGAATCAAGATAATAGACCAGTCACTATATTT |
| **Ribosome Binding Sites** | |
| B0032 | TCACACAGGAAAGTACTAG |
| $sRBS1_{mtrC}$[1] | GGGGAAAAACAGCAGTGCGAT |
| $sRBS0_{mtrA}$ [1] | ATAGGCGGCTTCATTGACGGTCCCA |
| $sRBS4_{cymA}$ [1] | TTCGCTTTGGGTTTTTAAGGAGGACGCA |
| tetR RBS[3] | CTATGGACTATGTTTTCACACAGGAAAGGCCTCG |
| lux1 RBS[5] | GGAAGAGAGTCAATTCAGGGTGGTGAAT |
| betI RBS [3] | GCTACGACTTGCTCATTTGACAGAGGATAACTACTA |
| qacR RBS [3] | GCCATGCCATTGGCTTTTGATAGAGGACAACTACTAG |
| phlF RBS [3] | CTATGGACTATGTTTGAAAGGGAGAAATACTAG |
| lmrA RBS [3] | CTATGGACTATGTTTTCACACAGGAAAGGGCTCG |
| **Terminators** | |
| T0 | CTTGGACTCCTGTTGATAGATCCAGTAATGACCTCAGAACTCCATCTGGATTTGTTCAGAACGCTCGGTTGCCGCCGGGCGTTTTTTATTGGTGAGAATCCAAGCA |
| ECK120029600[8] | TTCAGCCAAAAAACTTAAGACCGCCGGTCTTGTCCACTACCTTGCAGTAATGCGGTGGACAGGATCGGCGGTTTTCTTTTCTCTTCTCAA |
| L3S2P21[8] | TCGGTACCAAATTCCAGAAAAGAGGCCTCCCGAAAGGGGGGCCTTTTTTCGTTTTGGTCC |
| B0015 | TAATCTAGACCAGGCATCAAATAAAACGAAAGGCTCAGTCGAAAGACTGGGCCTTTCGTTTTATCTGTTGTTTGTCGGTGAACGCTCTCTACTAGAGTCACACTGGCTCACCTTCGGGTGGGCCTTTCTGCGTTTATA |
| ECK120033737[8] | GGAAACACAGAAAAAAGCCCGCACCTGACAGTGCGGGCTTTTTTTTTCGACCAAAGG |
| L3S2P55[8] | CTCGGTACCAAAGACGAACAATAAGACGCTGAAAAGCGTCTTTTTTCGTTTTGGTCC |
| ECK120033736[8] | AACGCATGAGAAAGCCCCCGGAAGATCACCTTCCGGGGGCTTTTTTATTGCGC |
| ECK120017009[8] | GATCTAACTAAAAAGGCCGCTCTGCGGCCTTTTTTCTTTTCACT |
| **Ribozymes** | |
| RiboJ[9] | AGCTGTCACCGGATGTGCTTTCCGGTCTGATGAGTCCGTGAGGACGAAACAGCCTCTACAAATAATTTTGTTTAA |

| | |
|---|---|
| ElvJ[9] | AGCCCCATAGGGTGGTGTGTACCACCCCTGATGAGTCCAAAAGGACGAAATGGGGCCTCTACAA<br>ATAATTTTGTTTAA |
| PlmJ[9] | AGTCATAAGTCTGGGCTAAGCCCACTGATGAGTCGCTGAAATGCGACGAAACTTATGACCTCTAC<br>AAATAATTTTGTTTAA |
| RiboJ64[9] | AGGAGTCAATTAATGTGCTTTTAATTCTGATGAGACGGTGACGTCGAAACTCCCTCTACAAATAAT<br>TTTGTTTAA |
| **Genes** | |
| *tetR*<br>(codon optimized for MR-1 using jcat.de) | ATGTCTCGTTTAGATAAATCTAAAGTTATCAACTCTGCTTTAGAATTATTAAACGAAGTTGGTATCG<br>AAGGTTTAACTACTCGTAAATTAGCTCAAAAATTAGGTGTTGAACAGCCCACATTATACTGGCACG<br>TTAAAAACAAGAGGGCGTTATTAGATGCTCTCGCTATCGAAATGTTAGATCGTCACCACACTCACT<br>TCTGTCCATTAGAAGGTGAATCTTGGCAAGATTTCTTACGTAACAACGCTAAATCGTTCCGTTGTG<br>CGTTATTATCGCACCGTGATGGTGCTAAAGTTCACTTAGGTACTCGTCCAACTGAAAAACAATACG<br>AAACTTTAGAAAACCAATTAGCTTTCTTATGTCAACAAGGTTTCTCGCTCGAAAACGCGCTCTATG<br>CGTTATCGGCTGTTGGCCACTTCACTTTAGGTTGTGTTTTAGAAGATCAAGAACACCAAGTTGCTA<br>AAGAAGAACGTGAAACTCCAACTACTGATTCTATGCCACCATTATTACGTCAAGCTATCGAATTAT<br>TCGATCACCAAGGTGCCGAGCCAGCGTTCCTCTTCGGTTTAGAATTAATCATCTGTGGTTTAGAA<br>AAACAATTAAAATGTGAATCTGGTTCTTAA |
| *lacI* | GTGAAACCAGTAACGTTATACGATGTCGCAGAGTATGCCGGTGTCTCTTATCAGACCGTTTCCCG<br>CGTGGTGAACCAGGCCAGCCACGTTTCTGCGAAAACGCGGGAAAAAGTGGAAGCGGCGATGGC<br>GGAGCTGAATTACATTCCCAACCGCGTGGCACAACAACTGGCGGGCAAACAGTCGTTGCTGATT<br>GGCGTTGCCACCTCCAGTCTGGCCCTGCACGCGCCGTCGCAAATTGTCGCGGCGATTAAATCTC<br>GCGCCGATCAACTGGGTGCCAGCGTGGTGGTGTCGATGGTAGAACGAAGCGGCGTCGAAGCCT<br>GTAAAGCGGCGGTGCACAATCTTCTCGCGCAACGCGTCAGTGGGCTGATCATTAACTATCCGCT<br>GGATGACCAGGATGCCATTGCTGTGGAAGCTGCCTGCACTAATGTTCCGGCGTTATTTCTTGATG<br>TCTCTGACCAGACACCCATCAACAGTATTATTTTCTCCCATGAAGACGGTACGCGACTGGGCGTG<br>GAGCATCTGGTCGCATTGGGTCACCAGCAAATCGCGCTGTTAGCGGGCCCATTAAGTTCTGTCT<br>CGGCGCGTCTGCGTCTGGCTGGCTGGCATAAATATCTCACTCGCAATCAAATTCAGCCGATAGC<br>GGAACGGGAAGGCGACTGGAGTGCCATGTCCGGTTTTCAACAAACCATGCAAATGCTGAATGAG<br>GGTATCGTTCCCACTGCGATGCTGGTTGCCAACGATCAGATGGCGCTGGGCGCAATGCGCGCC<br>ATTACCGAGTCCGGGCTGCGCGTTGGTGCGGATATCTCGGTAGTGGGATACGACGATACCGAAG<br>ACAGCTCATGTTATATCCCGCCGTTAACCACCATCAAACAGGATTTTCGCCTGCTGGGGCAAACC<br>AGCGTGGACCGCTTGCTGCAACTCTCTCAGGGCCAGGCGGTGAAGGGCAATCAGCTGTTGCCC<br>GTGTCACTGGTGAAAAGAAAAACCACCCTGGCGCCCAATACGCAAACCGCCTCTCCCCGCGCGT<br>TGGCCGATTCATTAATGCAGCTGGCACGACAGGTTTCCCGACTGGAAAGCGGGCAGTGA |
| *luxR* | ATGAAAAACATAAATGCCGACGACACATACAGAATAATTAATAAAATTAAAGCTTGTAGAAGCAAT<br>AATGATATTAATCAATGCTTATCTGATATGACTAAAATGGTACATTGTGAATATTATTTACTCGCGA<br>TCATTTATCCTCATTCTATGGTTAAATCTGATATTTCAATCCTAGATAATTACCCTAAAAAATGGAG<br>GCAATATTATGATGACGCTAATTTAATAAAATATGATCCTATAGTAGATTATTCTAACTCCAATCATT<br>CACCAATTAATTGGAATATATTTGAAAACAATGCTGTAAATAAAAAATCTCCAAATGTAATTAAAGA<br>AGCGAAAACATCAGGTCTTATCACTGGGTTTAGTTTCCCTATTCATACGGCTAACAATGGCTTCGG<br>AATGCTTAGTTTTGCACATTCAGAAAAAGACAACTATATAGATAGTTTATTTTTACATGCGTGTATG<br>AACATACCATTAATTGTTCCTTCTCTAGTTGATAATTATCGAAAAATAAATATAGCAAATAATAAATC<br>AAACAACGATTTAACCAAAAGAGAAAAAGAATGTTTAGCGTGGGCATGCGAAGGAAAAAGCTCTT<br>GGGATATTTCAAAAATATTAGGTTGCAGTGAGCGTACTGTCACTTTCCATTTAACCAATGCGCAAA<br>TGAAACTCAATACAACAAACCGCTGCCAAAGTATTTCTAAAGCAATTTTAACAGGAGCAATTGATT<br>GCCCATACTTTAAAAATTGA |
| *phlF*[3] | CTATGGACTATGTTTGAAAGGGAGAAATACTAGATGGCACGTACCCCGAGCCGTAGCAGCATTG<br>GTAGCCTGCGTAGTCCGCATACCCATAAAGCAATTCTGACCAGCACCATTGAAATCCTGAAAGAA<br>TGTGGTTATAGCGGTCTGAGCATTGAAAGCGTTGCACGTCGTGCCGGTGCAAGCAAACCGACCA<br>TTTATCGTTGGTGGACCAATAAAGCAGCACTGATTGCCGAAGTGTATGAAAATGAAAGCGAACAG<br>GTGCGTAAATTTCCGGATCTGGGTAGCTTTAAAGCCGATCTGGATTTTCTGCTGCGTAATCTGTG<br>GAAAGTTTGGCGTGAAACCATTTGTGGTGAAGCATTTCGTTGTGTTATTGCAGAAGCACAGCTGG<br>ACCCTGCAACCCTGACCCAGCTGAAAGATCAGTTTATGGAACGTCGTCGTGAGATGCCGAAAAAA<br>CTGGTTGAAAATGCCATTAGCAATGGTGAACTGCCGAAAGATACCAATCGTGAACTGCTGCTGGA<br>TATGATTTTTGGTTTTTGTTGGTATCGCCTGCTGACCGAACAGCTGACCGTTGAACAGGATATTGA<br>AGAATTTACCTTCCTGCTGATTAATGGTGTTTGTCCGGGTACACAGCGT |
| *betI*[3] | GTGCCGAAACTGGGTATGCAGAGCATTCGTCGTCGTCAGCTGATTGATGCAACCCTGGAAGCAA<br>TTAATGAAGTTGGTATGCATGATGCAACCATTGCACAGATTGCACGTCGTGCCGGTGTTAGCACC<br>GGTATTATTAGCCATTATTTCCGCGATAAAAACGGTCTGCTGGAAGCAACCATGCGTGATATTACC<br>AGCCAGCTGCGTGATGCAGTTCTGAATCGTCTGCATGCACTGCCGCAGGGTAGCGCAGAACAGC<br>GTCTGCAGGCAATTGTTGGTGGTAATTTTGATGAAACCCAGGTTAGCAGCGCAGCAATGAAAGCA<br>TGGCTGGCATTTTGGGCAAGCAGCATGCATCAGCCGATGCTGTATCGTCTGCAGCAGGTTAGCA<br>GTCGTCGTCTGCTGAGCAATCTGGTTAGCGAATTTCGTCGTGAACTGCCTCGTGAACAGGCACAA<br>GAGGCAGGTTATGGTCTGGCAGCACTGATTGATGGTCTGTGGCTGCGTGCAGCACTGAGCGGTA<br>AACCGCTGGATAAAACCCGTGCAAATAGCCTGACCCGTCATTTTATCACCCAGCATCTGCCGACC<br>GATTAA |

| | |
|---|---|
| *qacR*[3] | ATGAACCTGAAAGATAAAATTCTGGGCGTTGCCAAAGAACTGTTTATCAAAAATGGCTATAACGCA<br>ACCACCACCGGTGAAATTGTTAAACTGAGCGAAAGCAGCAAAGGCAATCTGTATTATCACTTTAAA<br>ACCAAAGAGAACCTGTTTCTGGAAATCCTGAACATCGAAGAAAGCAAATGGCAAGAGCAGTGGAA<br>AAAAGAACAAATCAAATGCAAAACCAACCGCGAGAAATTCTATCTGTATAATGAACTGAGCCTGAC<br>CACCGAATATTACTATCCGCTGCAGAATGCCATCATCGAGTTTTATACCGAGTACTATAAAACCAA<br>CAGCATCAACGAGAAAATGAACAAACTGGAAAACAAATACATCGATGCCTACCACGTGATCTTTAA<br>AGAAGGTAATCTGAACGGCGAATGGTGCATTAATGATGTTAATGCCGTGAGCAAAATTGCAGCAA<br>ATGCCGTTAATGGCATTGTTACCTTTACCCATGAGCAGAATATCAACGAACGCATTAAACTGATGA<br>ACAAATTCAGCCAGATCTTTCTGAATGGCCTGAGCAAATAA |
| *lmrA*[3] | ATGAGCTATGGTGATAGCCGTGAAAAAATTCTGAGCGCAGCAACCCGTCTGTTTCAGCTGCAGG<br>GTTATTATGGCACCGGTCTGAATCAGATTATCAAAGAAAGCGGTGCACCGAAAGGTAGCCTGTAT<br>TATCATTTTCCGGGTGGTAAAGAACAGCTGGCAATTGAAGCAGTGAACGAAATGAAAGAATATAT<br>CCGCCAGAAAATCGCCGATTGTATGGAAGCATGTACCGATCCGGCAGAAGGTATTCAGGCATTT<br>CTGAAAGAACTGAGCTGTCAGTTTAGCTGTACCGAAGATATTGAAGGTCTGCCGGTTGGTCTGCT<br>GGCAGCAGAAACCAGCCTGAAAAGCGAACCGCTGCGTGAAGCATGTCATGAAGCATATAAAGAA<br>TGGGCCAGCGTGTATGAAGAAAAACTGCGTCAGACCGGTTGTAGCGAAAGCCGTGCAAAAGAAG<br>CAAGCACCGTTGTTAATGCAATGATTGAAGGTGGTATTCTGCTGAGCCTGACCGCAAAAAATAGC<br>ACACCGCTGCTGCATATTAGCAGCTGTATTCCGGATCTGCTGAAACGTTAA |
| *sfgfp* | ATGCGTAAAGGCGAAGAGCTGTTCACTGGTGTCGTCCCTATTCTGGTGGAACTGGATGGTGATG<br>TCAACGGTCATAAGTTTTCCGTGCGTGGCGAGGGTGAAGGTGACGCAACTAATGGTAAACTGAC<br>GCTGAAGTTCATCTGTACTACTGGTAAACTGCCGGTACCTTGGCCGACTCTGGTAACGACGCTGA<br>CTTATGGTGTTCAGTGCTTTGCTCGTTATCCGGACCATATGAAGCAGCATGACTTCTTCAAGTCC<br>GCCATGCCGGAAGGCTATGTGCAGGAACGCACGATTTCCTTTAAGGATGACGGCACGTACAAAA<br>CGCGTGCGGAAGTGAAATTTGAAGGCGATACCCTGGTAAACCGCATTGAGCTGAAAGGCATTGA<br>CTTTAAAGAAGACGGCAATATCCTGGGCCATAAGCTGGAATACAATTTTAACAGCCACAATGTTTA<br>CATCACCGCCGATAAACAAAAAAATGGCATTAAAGCGAATTTTAAAATTCGCCACAACGTGGAGG<br>ATGGCAGCGTGCAGCTGGCTGATCACTACCAGCAAAACACTCCAATCGGTGATGGTCCTGTTCT<br>GCTGCCAGACAATCACTATCTGAGCACGCAAAGCGTTCTGTCTAAAGATCCGAACGAGAAACGC<br>GATCATATGGTTCTGCTGGAGTTCGTAACCGCAGCGGGCATCACGCATGGTATGGATGAACTGT<br>ACAAATGATGA |
| *eYFP* | ATGGTGAGCAAGGGCGAGGAGCTGTTCACCGGGGTGGTGCCCATCCTGGTCGAGCTGGACGGC<br>GACGTAAACGGCCACAAGTTCAGCGTGTCCGGCGAGGGCGAGGGCGATGCCACCTACGGCAAG<br>CTGACCCTGAAGTTCATCTGCACCACCGGCAAGCTGCCCGTGCCCTGGCCCACCCTCGTGACCA<br>CCTTCGGCTACGGCCTGCAATGCTTCGCCCGCTACCCCGACCACATGAAGCTGCACGACTTCTT<br>CAAGTCCGCCATGCCCGAAGGCTACGTCCAGGAGCGCACCATCTTCTTCAAGGACGACGGCAAC<br>TACAAGACCCGCGCCGAGGTGAAGTTCGAGGGCGACACCCTGGTGAACCGCATCGAGCTGAAG<br>GGCATCGACTTCAAGGAGGACGGCAACATCCTGGGGCACAAGCTGGAGTACAACTACAACAGCC<br>ACAACGTCTATATCATGGCCGACAAGCAGAAGAACGGCATCAAGGTGAACTTCAAGATCCGCCA<br>CAACATCGAGGACGGCAGCGTGCAGCTCGCCGACCACTACCAGCAGAACACCCCCATCGGCGA<br>CGGCCCCGTGCTGCTGCCCGACAACCACTACCTGAGCTACCAGTCCGCCCTGAGCAAAGACCC<br>CAACGAGAAGCGCGATCACATGGTCCTGCTGGAGTTCGTGACCGCCGCCGGGATCACTCTCGG<br>CATGGACGAGCTGTACAAGTAA |
| *mtrC* | ATGATGAACGCACAAAAATCAAAAATCGCACTGCTGCTCGCAGCAAGTGCCGTCACAATGGCCTT<br>AACCGGCTGTGGTGGAAGCGATGGTAATAACGGCAATGATGGTAGTGATGGTGGTGAGCCAGCA<br>GGTAGCATCCAGACGTTAAACCTAGATATCACTAAAGTAAGCTATGAAAATGGTGCACCTATGGT<br>CACTGTTTTCGCCACTAACGAAGCCGACATGCCAGTGATTGGTCTCGCAAATTTAGAAATCAAAA<br>AAGCACTGCAATTAATACCGGAAGGGGCGACAGGCCCAGGTAATAGCGCTAACTGGCAAGGCTT<br>AGGCTCATCAAAGAGCTATGTCGATAATAAAAACGGTAGCTATACCTTTAAATTCGACGCCTTCGA<br>TAGTAATAAGGTCTTTAATGCTCAATTAACGCAACGCTTTAACGTTGTTTCTGCTGCGGGTAAATT<br>AGCAGACGGAACGACCGTTCCCGTTGCCGAAATGGTTGAAGATTTCGACGGCCAAGGTAATGCG<br>CCGCAATATACAAAAAATATCGTTAGCCACGAAGTATGTGCTTCTTGCCACGTAGAAGGTGAAAA<br>GATTTATCACCAAGCTACTGAAGTCGAAACTTGTATTTCTTGCCACACTCAAGAGTTTGCGGATGG<br>TCGCGGCAAACCCCATGTCGCCTTTAGTCACTTAATTCACAATGTGCATAATGCCAACAAAGCTT<br>GGGGCAAAGACAATAAAATCCCTACAGTTGCACAAAATATTGTCCAAGATAATTGCCAAGTTTGTC<br>ACGTTGAATCCGACATGCTCACCGAGGCAAAAAACTGGTCACGTATTCCAACAATGGAAGTCTGT<br>TCTAGCTGTCACGTAGACATCGATTTTGCTGCGGGTAAAGGCCACTCTCAACAACTCGATAACTC<br>CAACTGTATCGCCTGCCATAACAGCGACTGGACTGCTGAGTTACACACAGCCAAAACCACCGCA<br>ACTAAGAACTTGATTAATCAATACGGTATCGAGACTACCTCGACAATTAATACCGAAACTAAAGCA<br>GCCACAATTAGTGTTCAAGTTGTAGATGCGAACGGTACTGCTGTTGATCTCAAGACCATCCTGCC<br>TAAAGTGCAACGCTTAGAGATCATCACCAACGTTGGTCCTAATAATGCAACCTTAGGTTATAGTGG<br>CAAAGATTCAATATTTGCAATCAAAAATGGAGCTCTTGATCCAAAAGCTACTATCAATGATGCTGG<br>CAAACTGGTTTATACCACTACTAAAGACCTCAAACTTGGCCAAAACGGCGCAGACAGCGACACAG<br>CATTTAGCTTTGTAGGTTGGTCAATGTGTTCTAGCGAAGGTAAGTTTGTAGACTGTGCAGACCCT<br>GCATTTGATGGTGTTGATGTAACTAAGTATACCGGCATGAAAGCGGATTTAGCCTTTGCTACTTTG<br>TCAGGTAAAGCACCAAGTACTCGCCACGTTGATTCTGTTAACATGACAGCCTGTGCCAATTGCCA<br>CACTGCTGAGTTCGAAATTCACAAAGGCAAACAACATGCAGGCTTTGTGATGACAGAGCAACTAT<br>CACACACCCAAGATGCTAACGGTAAAGCGATTGTAGGCCTTGACGCATGTGTGACTTGTCATACT |

| | |
|---|---|
| | CCTGATGGCACCTATAGCTTTGCCAACCGTGGTGCGCTAGAGCTAAAACTACACAAAAAACACGT<br>TGAAGATGCCTACGGCCTCATTGGTGGCAATTGTGCCTCTTGTCACTCAGACTTCAACCTTGAGT<br>CTTTCAAGAAGAAAGGCGCATTGAATACTGCCGCTGCAGCAGATAAAACAGGTCTATATTCTACG<br>CCGATCACTGCAACTTGTACTACCTGTCACACAGTTGGCAGCCAGTACATGGTCCATACGAAAGA<br>AACCCTGGAGTCTTTCGGTGCAGTTGTTGATGGCACAAAAGATGATGCTACCAGTGCGGCACAG<br>TCAGAAACCTGTTTCTACTGCCATACCCCAACAGTTGCAGATCACACTAAAGTGAAAATGTAA |
| *mtrA* | ATGAAGAACTGCCTAAAAATGAAAAACCTACTGCCGGCACTTACCATCACAATGGCAATGTCTGC<br>AGTTATGGCATTAGTCGTCACACCAAACGCTTATGCGTCGAAGTGGGATGAGAAAATGACGCCAG<br>AGCAAGTCGAAGCCACCTTAGATAAGAAGTTTGCCGAAGGCAACTACTCCCCTAAAGGCGCCGA<br>TTCTTGCTTGATGTGCCATAAGAAATCCGAAAAAGTCATGGACCTTTTCAAAGGTGTCCACGGTG<br>CGATTGACTCCTCTAAGAGTCCAATGGCTGGCCTGCAATGTGAGGCATGCCACGGCCCACTGGG<br>TCAGCACAACAAAGGCGGCAACGAGCCGATGATCACTTTTGGTAAGCAATCAACCTTAAGTGCCG<br>ACAAGCAAAACAGCGTATGTATGAGCTGTCACCAAGACGATAAGCGTATGTCTTGGAATGGCGGT<br>CACCATGACAATGCCGATGTTGCTTGTGCTTCTTGTCACCAAGTACACGTCGCAAAAGATCCTGT<br>GTTATCTAAAAACACGGAAATGGAAGTCTGTACTAGCTGCCATACAAAGCAAAAAGCGGATATGA<br>ATAAACGCTCAAGTCACCCACTCAAATGGGCACAAATGACCTGTAGCGACTGTCACAATCCCCAT<br>GGGAGCATGACAGATTCCGATCTTAACAAGCCTAGCGTGAATGATACCTGTTATTCCTGTCACGC<br>CGAAAAACGCGGCCCAAAACTTTGGGAGCATGCACCCGTCACTGAGAATTGTGTCACTTGCCAC<br>AATCCTCACGGTAGTGTGAATGACGGTATGCTGAAAACCCGTGCGCCACAGCTATGTCAGCAAT<br>GTCACGCCAGCGATGGCCACGCCAGCAACGCCTACTTAGGTAACACTGGATTAGGTTCAAATGT<br>CGGTGACAATGCCTTTACTGGTGGAAGAAGCTGCTTAAATTGCCATAGTCAGGTTCATGGTTCTA<br>ACCATCCATCTGGCAAGCTATTACAGCGCTAA |
| *cymA* | ATGAACTGGCGTGCACTATTTAAACCCAGCGCGAAATATTCCATCCTAGCGCTACTGGTTGTTGG<br>TATCGTGATTGGTGTTGTGGGCTATTTTGCAACTCAGCAGACTTTACATGCGACAAGTACAGATG<br>CGTTCTGTATGTCTTGCCATAGCAATCATTCCTTGAAGAATGAAGTGCTGGCATCTGCCCACGGT<br>GGCGGCAAAGCCGGGGTTACTGTTCAGTGTCAAGACTGTCACTTACCCCATGGCCCTGTTGATT<br>ATTTAATTAAGAAAATCATCGTATCTAAAGATTTATATGGTTTCTTAACTATTGATGGCTTTAACACT<br>CAAGCTTGGTTAGACGAAAACCGCAAAGAGCAAGCCGACAAAGCATTGGCTTACTTCCGTGGTAA<br>CGACTCAGCAAACTGTCAACACTGCCATACTCGCATTTATGAAAACCAGCCAGAAACCATGAAGC<br>CAATGGCTGTGAGAATGCACACCAACAACTTCAAGAAAGATCCTGAAACGAGAAAGACCTGTGTG<br>GATTGCCACAAAGGTGTCGCTCACCCCTATCCAAAAGGATAA |

**Table S3**. Nonlinear (sigmoidal) gene expression fit parameters for products of *sfgfp* expression (fluorescence) or EET gene expression (storage modulus).

Parameters were fit using an activating or deactivating 4-parameter sigmoidal model (Hill Function) in Prism 9.

| Plasmid | GOI (condition) | *n* | $K_{1/2}$ (µM) | $y_{max}$ (units) | $y_{min}$ (units) | Dynamic Range | Goodness of Fit ($R^2$) |
|---|---|---|---|---|---|---|---|
| pCD7sfGFP | *sfgfp* | 3.4 | 280 | 0.17 (REU) | 0.021 (REU) | 8.1 | 0.76 |
| pCD24r1 | *mtrC* (stationary phase) | 1.1 | 32 | 400 (Pa) | 16 (Pa) | 25 | 0.47 |
| pCD24r1 | *mtrC* (dynamic) | 4.5 | 83 | 900 (Pa) | 37 (Pa) | 24 | 0.74 |
| pCD25r0 | *mtrA* (stationary phase) | 0.95 | 220 | 1000 (Pa) | 16 (Pa) | 61 | 0.60 |
| pCD25r0 | *mtrA* (dynamic) | 1.2 | 300 | 510 (Pa) | 15 (Pa) | 34 | 0.78 |
| pCD26r4 | *cymA* (stationary phase) | 2.1 | 22 | 170 (Pa) | 2.6 (Pa) | 65 | 0.69 |
| pCD26r4 | *cymA* (dynamic) | 1.2 | 81 | 160 (Pa) | 15 (Pa) | 11 | 0.69 |
| pAT3 | *sfgfp* | 1.1 | 0.017 | 0.63 (REU) | 0.0029 (REU) | 210 | 0.95 |
| pAT1 | *mtrC* (dynamic) | 2.4 | 0.030 | 700 (Pa) | 20 (Pa) | 35 | 0.72 |
| pCDTetRsfGFP | *sfgfp* | 2.1 | 0.0014 | 0.43 (REU) | 0.17 (REU) | 2.6 | 0.94 |
| pCD24tr1* | *mtrC* (dynamic) | 1.2 | 0.0010 | 380 (Pa) | 180 (Pa) | 2.1 | 0.21 |
| pCDd1 | *mtrC* (dynamic) | -0.91 | 10 | 2000 (Pa) | 33 (Pa) | 62 | 0.90 |
| pCD24r1 | *mtrC* (dynamic, CuAAC, THPTA) | 1.3 | 98 | 3800 (Pa) | 860 (Pa) | 4.4 | 0.66 |
| pCDd1 | *mtrC* (dynamic, CuAAC, THPTA) | -0.76 | 39 | 3000 (Pa) | 490 (Pa) | 6.0 | 0.79 |
| pCD24r1 | *mtrC* (dynamic, CuAAC, BTTAA) | 2.3 | 120 | 1600 (Pa) | 35 (Pa) | 45 | 0.77 |
| pCDd1 | *mtrC* (dynamic, CuAAC, BTTAA) | -17 | 53 | 1100 (Pa) | 7.5 (Pa) | 700 | 0.66 |

*Average storage modulus at no induction (0 nM aTc) was excluded from this fit as an outlier.

**Table S4**. Read-only Benchling links for new architectures used in this study, including primer and sequencing information.

| Plasmids | Description | Read-Only Benchling Link |
|---|---|---|
| pCDe1 | *sfgfp* constitutive REU (Ptrc* promoter) | pCDe1 |
| pCD24tr1 | *mtrC* Buffer gate ($P_{Tet}$; *tetR*) | pCD24r1 |
| pCDTetRsfGFP | *sfgfp* Buffer gate ($P_{Tet}$; *tetR*) | pCDTetRsfGFP |
| pAT1 | *mtrC* Buffer gate ($P_{Lux}$; *luxR*) | pAT1 |
| pAT3 | *sfgfp* Buffer gate ($P_{Lux}$; *luxR*) | pAT3 |
| pAJGb1-*mtrC* | *mtrC* OR gate (($P_{Tet}$; *tetR*; $P_{Lux}$; *luxR*) | pAJGb1-*mtrC* (OR) |
| pAJGb1-*sfgfp* | *sfgfp* OR gate (($P_{Tet}$; *tetR*; $P_{Lux}$; *luxR*) | pAJGb1-*sfgfp* (OR) |
| pAJGb2-*mtrC* | *mtrC* NOR gate (($P_{Tet}$; *tetR*; $P_{Lux}$; *luxR*) | pAJGb2-*mtrC* (NOR) |
| pAJGb2-*sfgfp* | *sfgfp* NOR gate (($P_{Tet}$; *tetR*; $P_{Lux}$; *luxR*) | pAJGb2-*sfgfp* (NOR) |
| pAJGb3-*mtrC* | *mtrC* AND gate ($P_{tacsym0}$; *lacI*; $P_{Tet}$; *tetR*) | pAJGb3-*mtrC* (AND); [3] |
| pAJGb3-*eYFP* | *eYFP* AND gate ($P_{tacsym0}$; *lacI*; $P_{Tet}$; *tetR*) | pAJGb3-*eYFP* (AND); [3] |
| pAJGb4-*mtrC* | *mtrC* NAND gate ($P_{tacsym0}$; *lacI*; $P_{Lux}$; *luxR*) | pAJGb4-*mtrC* (NAND); [3] |
| pAJGb4-*sfgfp* | *sfgfp* NAND gate ($P_{tacsym0}$; *lacI*; $P_{Lux}$; *luxR*) | pAJGb4-*sfgfp* (NAND); [3] |

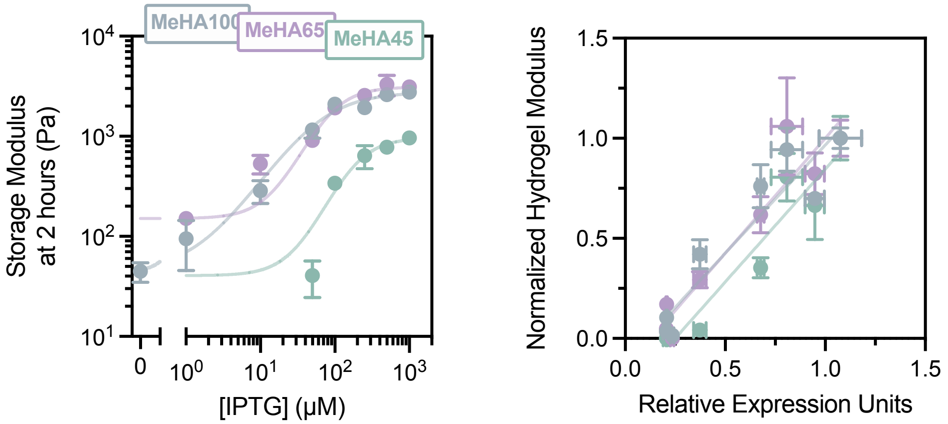

**Figure S1.** Storage modulus is transcriptionally regulated independent of methacrylate functionalization.

(Left) Methacrylated hyaluronic acid at different functional densities (45%, 65%, and 100% as measured by $^1$H-NMR[10]) was cross-linked using *S. oneidensis* Δ*mtrC*Δ*omcA*Δ*mtrF* + *mtrC* under LacI regulation at stationary phase. (Right) Storage modulus normalized to gels cross-linked using an empty vector control plotted as a function of relative gene expression determined using a fluorescent circuit under identical transcriptional regulation. Data shown are mean ± SEM of *n* = 3 biological replicates.

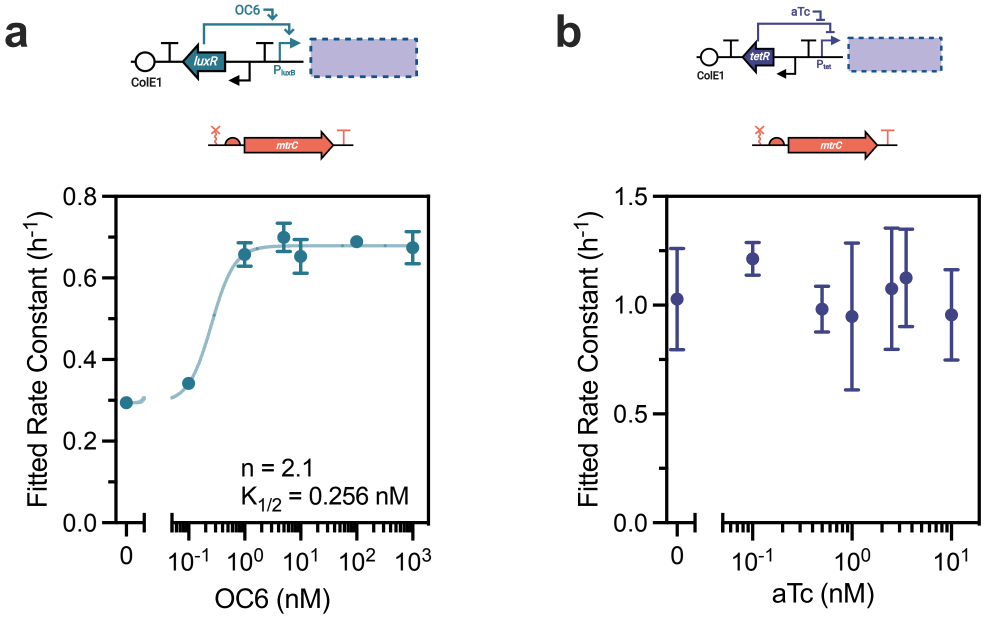

**Figure S2**. Iron reduction rate constant response functions for LuxR- and TetR-regulated Buffer gates controlling *mtrC* expression.

*In situ* Fe(III) reduction data are fit to a Monod-type model as described in the Methods to obtain fitted rate constants; the **a**, LuxR-regulated rate constants are then fit to a sigmoidal gene expression model. The **b**, TetR-regulated circuit was not fit to a gene expression model as it did not exhibit sigmoidal behavior. Data shown are mean ± SEM of *n* = 3 biological replicates.

**Table S5.** Plasmid maps of new architectures used in this study.

Below each map a read-only Benchling link is provided for the plasmid, including sequencing and primer information.

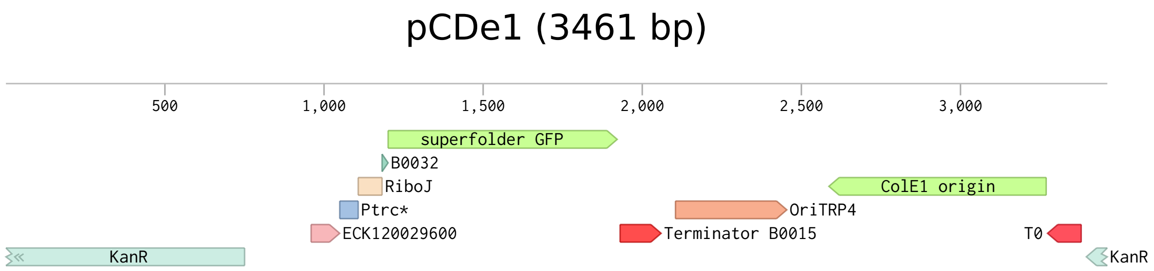

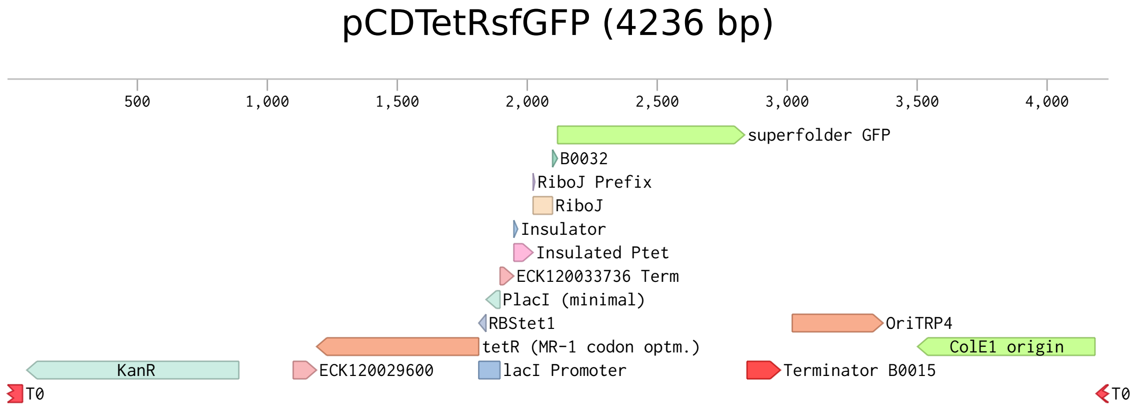

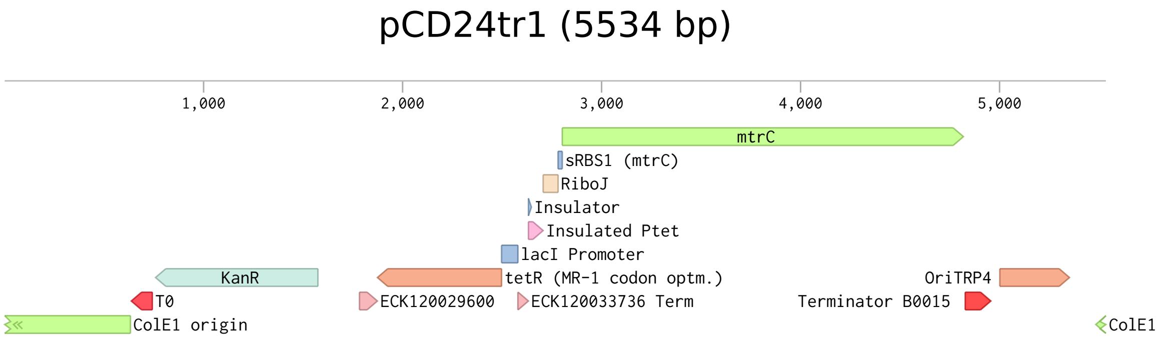

#### pAT3 (4337 bp)

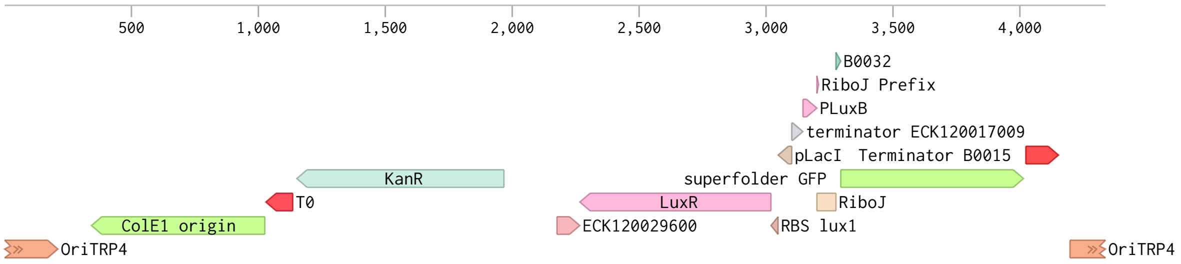

#### pAT1 (5635 bp)

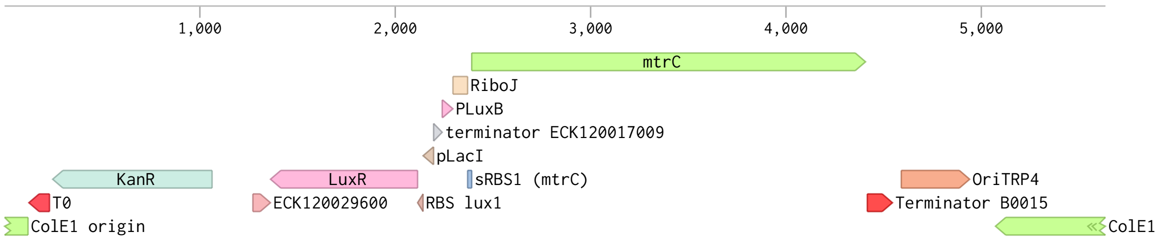

#### pAJGb1-sfgfp (OR) (5108 bp)

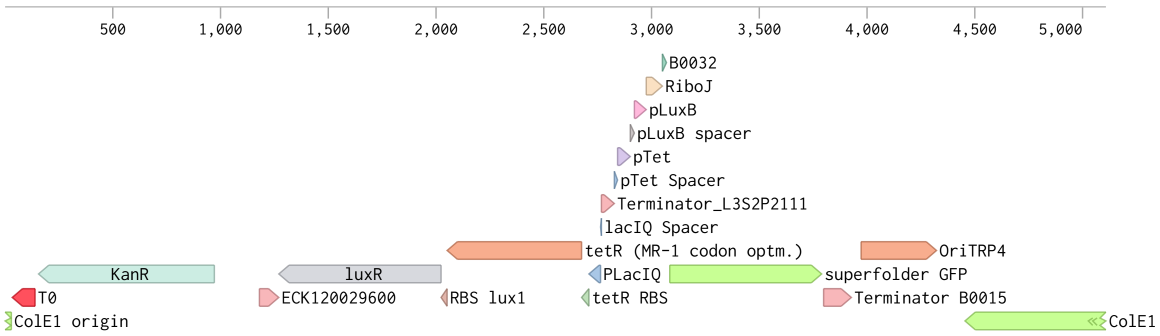

### pAJGb1-mtrC (OR) (6406 bp)

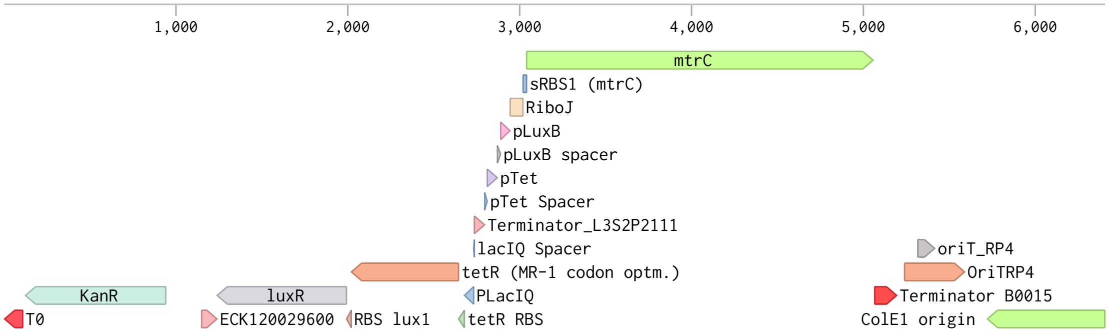

### pAJGb2-sfgfp II (PhlF Design) (5999 bp)

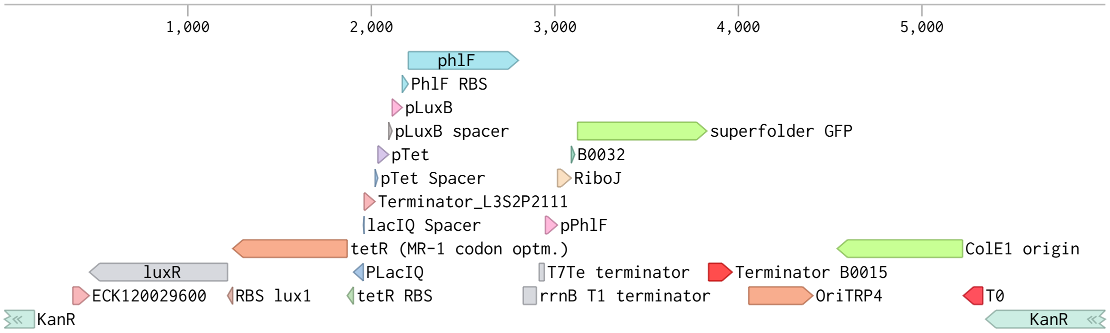

### pAJGb2-mtrC II (PhlF Design) (7297 bp)

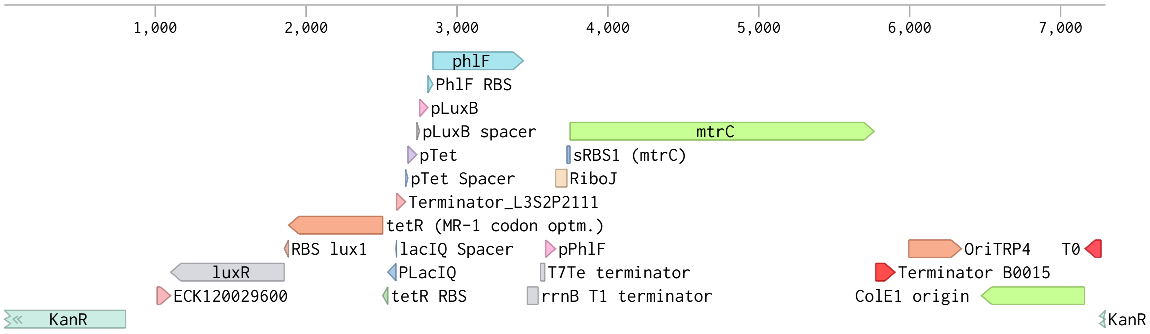

### pAJGb3-eYFP (AND) (8270 bp)

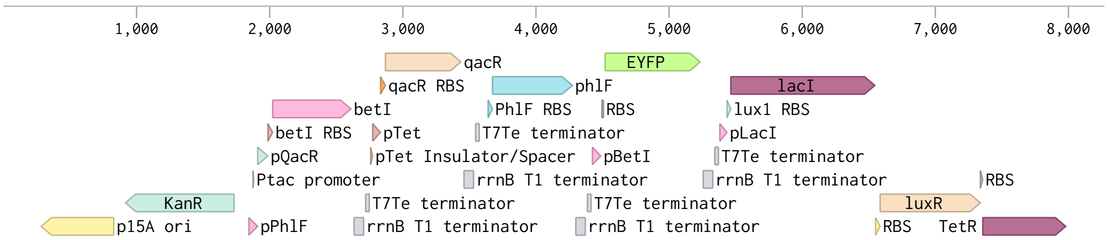

### pAJGb3-mtrC (AND) (9681 bp)

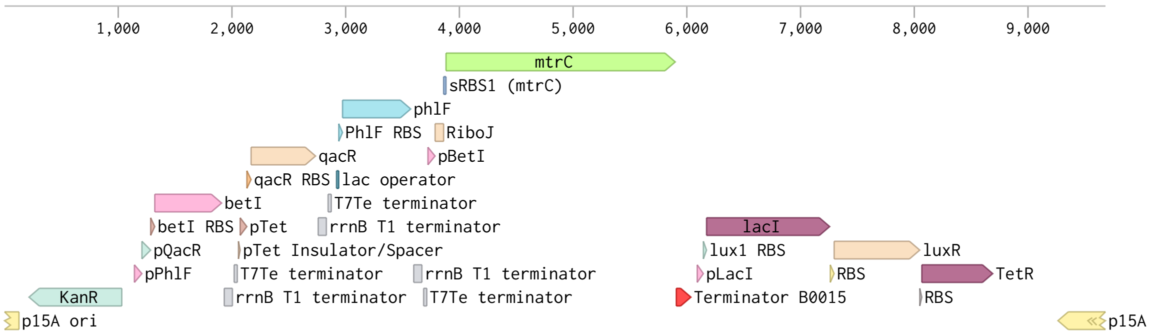

### pAJGb4-sfgfp (NAND) (7261 bp)

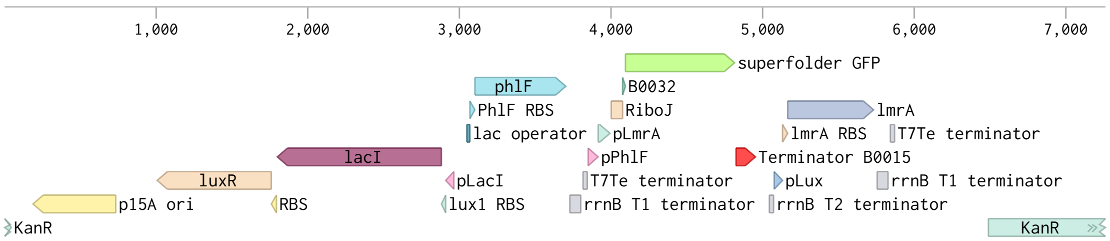

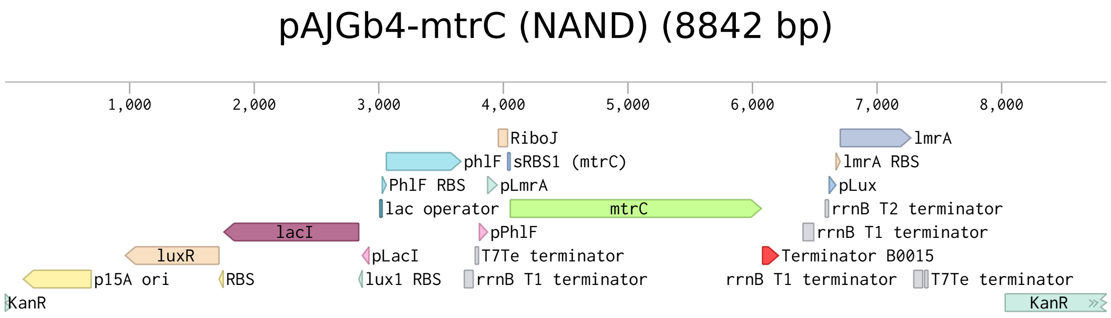
pAJGb4-mtrC (NAND) (8842 bp)
1,000
2,000
3,000
4,000
5,000
6,000
7,000
8,000
RiboJ
lmrA
phlF
sRBS1 (mtrC)
lmrA RBS
PhlF RBS
pLmrA
pLux
lac operator
mtrC
rrnB T2 terminator
lacI
pPhlF
rrnB T1 terminator
luxR
pLacI
T7Te terminator
Terminator B0015
p15A ori
RBS
lux1 RBS
rrnB T1 terminator
rrnB T1 terminator
T7Te terminator
KanR
KanR

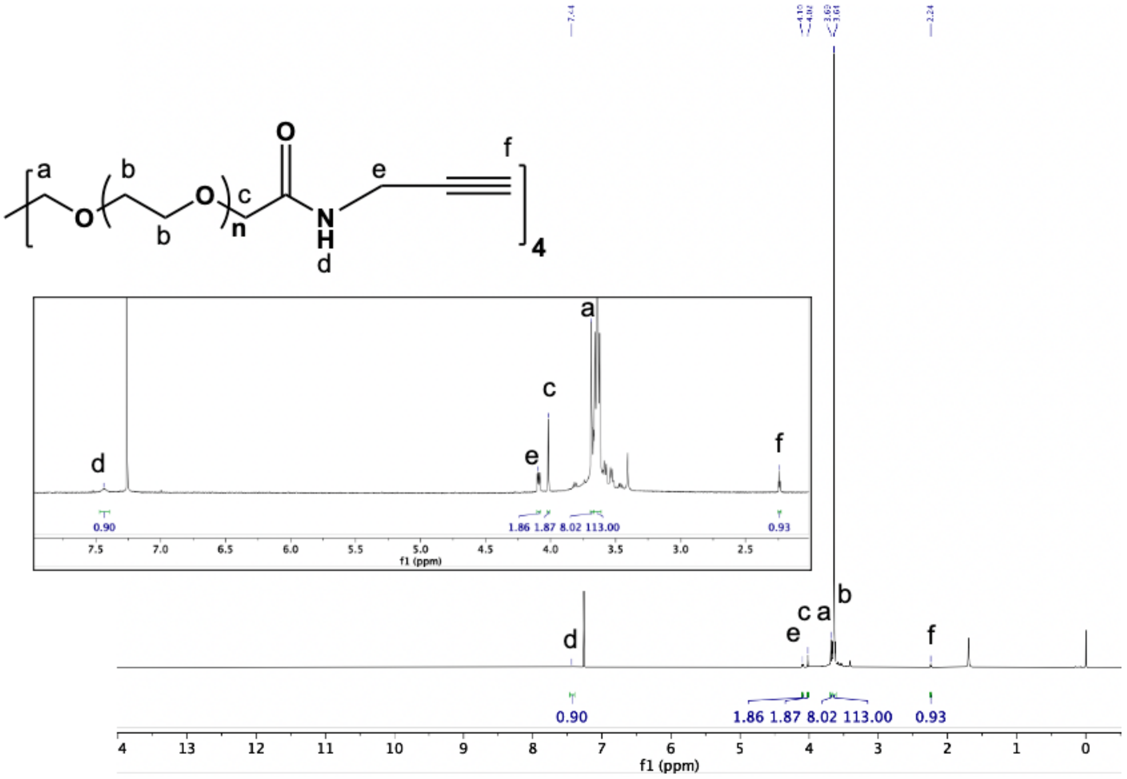

**Figure S3.** $^{1}$H-NMR of synthesized PEG-Alkyne (see Methods).
